# Artificial DNA-nano/microparticle motors: Factors governing speed, run-length, and unidirectionality revealed by geometry-based kinetic simulations

**DOI:** 10.64898/2026.02.11.705463

**Authors:** Takanori Harashima, Ryota Iino

## Abstract

DNA-nano/microparticle motors are burnt-bridge Brownian ratchets (BBR) moving on an RNA-modified two-dimensional surface driven by Ribonuclease H (RNase H), and are one of the fastest artificial molecular motors. Interestingly, these motors show a maximum speed of ∼30 nm s⁻¹ irrespective of the particle size ranging from 100 to 5000 nm, whereas the run-length increases with the particle size. Here we performed geometry-based kinetic simulations of DNA-nano/microparticle motors with the sizes of 100, 500, 1000, and 5000 nm to identify the factors governing speed, run-length, and unidirectionality. The simulations reproduced the experiments quantitatively, and the speed remained constant while the run-length and the unidirectionality increased with the particle size. The constant speed was caused by a trade-off between the step size and the pause length, both of which increased with the particle size. In contrast, the run-length and the unidirectionality increased with the particle size because large particles had high multivalency which suppresses stochastic detachment of the motor, high RNA hydrolysis efficiency under the motor trajectory which realizes almost perfect BBR motion, and stepping direction highly biased to forward. For the smaller motors with 100, 500, and 1000 nm particles, the speed increased from 20 to 200 nm s⁻¹ by 10-fold increases in DNA/RNA hybridization, RNase H binding, and RNA hydrolysis rates (from 0.8 to 8.0, 7.2 to 72, and 3.0 to 30 s⁻¹, respectively), even when considering the rotational diffusion of these particles. On the other hand, the speed for the largest motor with 5000 nm particle was limited to 100 nm s⁻¹, because the time required for rolling motion (∼0.3 s) became comparable to the pause length. Our results indicate that DNA-particle motors must possess a nanoscale body to achieve a speed exceeding 100 nm s⁻¹.

**Significance:** Autonomous artificial molecular motors have a potential to power nano- and micron-scale actuators and devices, but their performances such as speed, run-length, and unidirectionality are inferior to natural motor proteins. Using geometry-based kinetic simulations, we quantitatively analyzed performance metrics of artificial DNA-nano/microparticle motors which autonomously move on RNA-modified two-dimensional surfaces by a burnt-bridge Brownian ratchet mechanism. Our study revealed the mechanism why their speed is almost independent of the particle size, while the run-length and unidirectionality increases with the particle size. We also identified how multivalent binding, mode of detachment, and rotational diffusion set fundamental limits of the speed, run-length, and unidirectionality. Our results provide a general design strategy for engineering high-performance artificial molecular motors.

## Introduction

Motor proteins such as myosin and kinesin convert chemical energy into mechanical work and play essential roles in activities of the living organisms such as muscle contraction and intracellular cargo transport (1–5). Their ability to rectify Brownian motion into autonomous unidirectional motion has inspired extensive efforts to construct artificial molecular motors that mimic or even rival biological ones (6). DNA walkers represent pioneering examples of artificial molecular motors that achieve nanoscale unidirectional motions (7,8). Some DNA walkers move along nucleic acid rails through successive hybridization and subsequent cleavage reactions, rectifying Brownian motion based on the burnt-bridge Brownian ratchet (BBR) mechanism. In this process, the DNA walker binds to RNA (or DNA) footholds and enzymatically cleaves (or “burns”) them, biasing the Brownian motion toward the intact footholds. Although their motions are autonomous, the speed is only ∼0.1 nm s⁻^1^ (7,9,10), which is much lower than that of motor proteins (typically 10–1000 nm s⁻^1^) (11–14). This large speed gap has been one of the central challenges in realizing artificial molecular motors that approach the performance of biological ones.

Recent developments of Ribonuclease H (RNase H)-driven DNA-nano/microparticle motors have demonstrated that multivalent BBR mechanisms can realize performances comparable to the biological motor proteins (15). DNA-modified microparticle motors with a diameter of 5 μm (5000 nm) moved autonomously on the RNA-modified surfaces via enzymatic RNA hydrolysis and showed a mean speed of ∼30 nm s⁻^1^ at high RNase H concentration ([RNase H]), which is comparable to that of a BBR-based motor protein (Table 1) (14,16). In the same report, submicron 500-nm particle motors also showed similar ∼30 nm s⁻^1^ speed. More recently, a nanoparticle motor with a diameter of 50 nm showed instantaneous maximum speed of 50 nm s⁻^1^ (17), and that with a diameter of 100 nm showed a mean speed of ∼30 nm s⁻^1^ (Table 1) (18). These studies establish DNA-nano/microparticle motors as one of the fastest artificial molecular motors reported to date. In parallel, another high-speed BBR motor, the “Lawnmower”, has been reported recently (19). In this artificial molecular motor, protease-modified particle with a diameter of 2.8 μm moved autonomously by degrading substrate peptides attached to the surface with a mean speed of 50–60 nm s⁻^1^, supporting the generality of BBR mechanisms across different molecular systems.

**Table 1.**
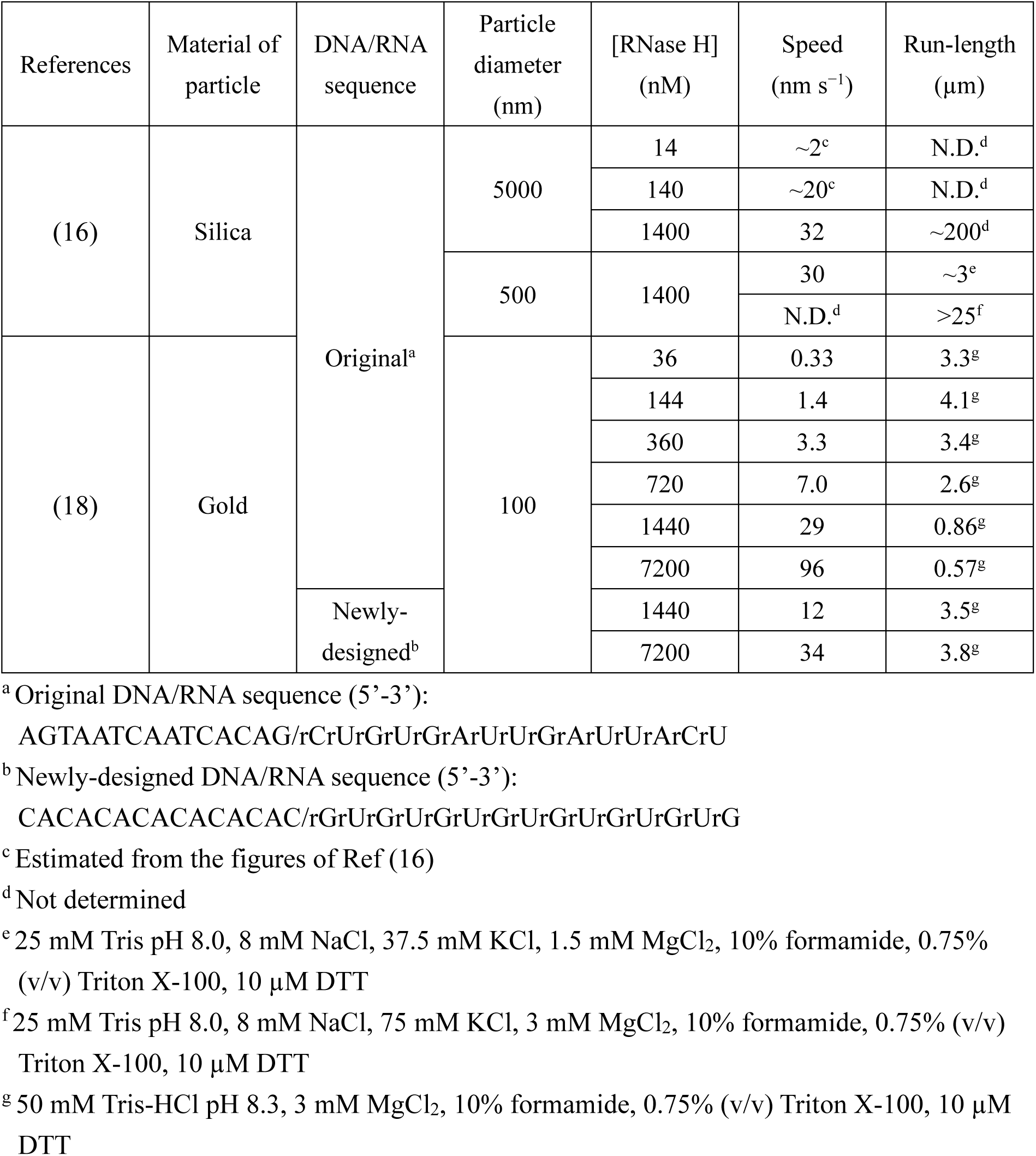
Performance metrics of the DNA-nano/microparticle motors with different DNA/RNA sequences and particle sizes.

An intriguing feature of DNA-nano/microparticle motors is that despite the largely different sizes of the motor bodies, the same BBR mechanism is at work. Furthermore, interestingly, the speed at high [RNase H] remains nearly constant (∼30 nm s⁻¹) over more than one order of difference in the particle diameter (100–5000 nm), indicating a scale-free property (Table 1) (16,18). In contrast, other performance metrics such as run-length and processivity increase with the particle diameter, indicating that not all performance metrics are scale-free. Despite extensive experimental characterizations, mechanistic origin of the coexistence of the size-independent speed and the size-dependent run-length and processivity remains elusive.

In parallel with the observations above, considerable effort has been devoted to improving motor performance through optimization of the DNA/RNA sequences. For the microparticle motor, variations in the GC content have been shown to modulate the speed, demonstrating that hybridization kinetics directly affects the motility (20). For the nanoparticle motor, recent experiments revealed a trade-off between the speed and the run-length (or processivity), which was successfully improved by introducing a repetitive GC-rich sequence to increase DNA/RNA hybridization rate (18). These studies demonstrated DNA/RNA sequence engineering as an effective approach to improve motor performance.

As another performance metric of the DNA-nano/microparticle motors, attention has also been focused on the unidirectionality, because they move on two-dimensional RNA-modified surface with the BBR mechanism in which only backward motion is suppressed but forward and side motions are allowed. In this aspect, analysis of step angle introduced in the previous study for 100-nm DNA-nanoparticle motor enabled quantitative evaluation of the unidirectionality at the level of elementary processes of the motion (pauses and steps), and revealed that DNA/RNA sequence engineering improved not only the speed and the run-length but also the unidirectionality at high [RNase H] (18). Recently, similar analysis of the unidirectionality has also been applied to the microparticle motors (21).

The particle size dependence described above suggests that the performances of DNA-nano/microparticle motors are governed by a delicate interplay among multivalency, kinetics of chemical reaction, and diffusion process. However, it has not been systematically investigated how the particle size affects their speed, run-length, and unidirectionality. Although previous theoretical studies have examined individual aspects such as substrate (foothold) stiffness (22) and multivalent binding/unbinding kinetics (23), a unified mechanism that explains the particle size dependence is still missing.

In the present study, we address this issue by systematically applying the geometry-based kinetic simulations (18) to the DNA-nano/microparticle motors with different particle diameters. Our simulations quantitatively reproduce the experimentally observed behaviors: the nearly constant speed (∼30 nm s⁻^1^) across the particle diameters from 100 to 5000 nm and the run-length and processivity increasing with the particle diameter. We show that particle size-independent speed arises from a compensation between the step size and the pause length, both of which increase with the particle diameter. In contrast, the increase in the run-length and processivity with the particle diameter is explained by a shift from stochastic detachment (complete hydrolysis of DNA/RNA duplexes under the motor before formation of new DNA/RNA duplexes) to entrap detachment (entrapment of the motor within a region where surrounding RNAs are already hydrolyzed) (18), caused by increased multivalency. Furthermore, we reveal an origin of the improved unidirectionality of the micron-sized motors. In short, more efficient RNA hydrolysis under the motor trajectory, together with statistical averaging over a large number of DNA/RNA duplexes due to increased multivalency, results in stepping direction highly biased to forward. Finally, by systematically varying the kinetic parameters of the elementary processes of the chemical reaction, we identify conditions in which the nano-sized motor achieves ∼200 nm s⁻^1^ speed and micrometer run-length simultaneously. Importantly, we also show that the speed of micron-sized motor becomes limited by the rotational diffusion under such fast reaction kinetics. Collectively, our results reveal the pros and cons of the nano- and micron-sized DNA-particle motors and provide a general design strategy to further enhance their performance metrics.

## Materials and Methods

### Model and algorithm of geometry-based kinetic simulation

The geometry-based kinetic simulations of DNA-nano/microparticle motors were implemented in Python 3.9. The model describes a spherical particle modified with single-stranded DNAs (ssDNA) moving on a surface modified with single-stranded RNAs, driven by RNase H-catalyzed RNA hydrolysis (Fig. 1a). The RNA surface was represented as a two-dimensional pixel matrix, where each pixel corresponds to a potential RNA site (foothold). Pixel size was normalized by the DNA density on the particle (0.10 molecules nm⁻^2^), as determined in our previous work (18). RNAs were then distributed by randomly selecting a fraction of pixels corresponding to the experimentally measured RNA density (0.022 molecules nm⁻^2^) on the surface. All geometric and kinetic parameters were chosen to match the conditions of our previous experiments and simulations on 100-nm DNA-gold nanoparticle motor (18) and are summarized in Table S1.

**Figure 1.**
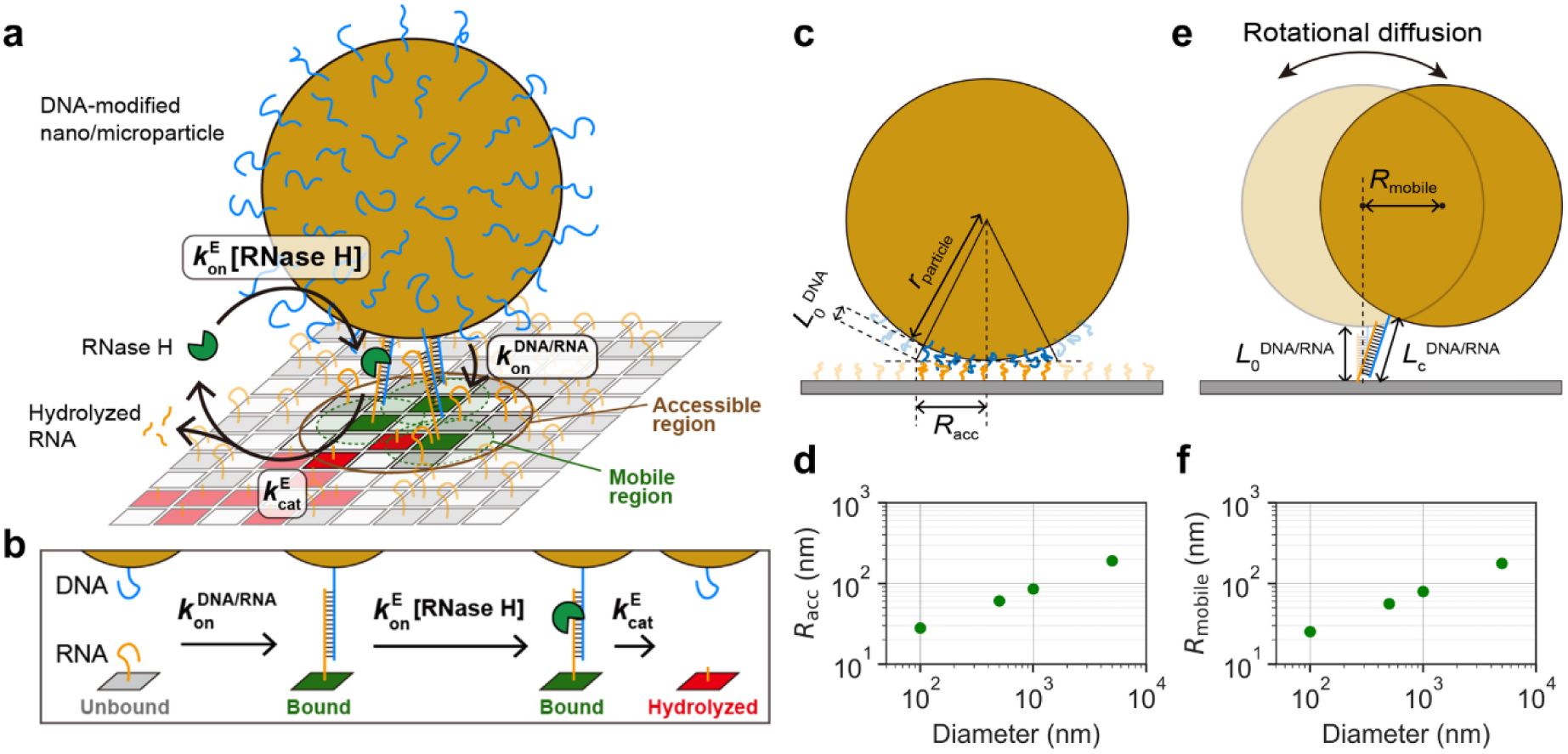
Framework of geometry-based kinetic simulation of DNA-nano/microparticle motors. (a) Schematic illustration of a DNA-modified nano/microparticle interacting with RNA-modified two-dimensional surface. Single-stranded DNAs on the particle hybridize with complementary single-stranded RNAs on the surface, RNase H enzymes bind to the formed DNA/RNA duplexes and selectively hydrolyze RNAs, and generate hydrolyzed RNA sites where DNA cannot bind again. (b) Reaction scheme consisting of three chemical processes: DNA/RNA hybridization with the rate of *k*_on_^DNA/RNA^, RNase H binding to DNA/RNA duplex with the rate of *k*_on_^E^[RNase H], and RNA hydrolysis and duplex dissociation with the rate of *k*_cat_^E^. These kinetic parameters were used as inputs to the Gillespie algorithm. (c) Geometric model used to estimate the accessible region, defined as the area in which DNAs on the particle can interact with RNAs on the surface. (d) Calculated accessible radius (*R*_acc_) for the particles with diameters of 100, 500, 1000, and 5000 nm. (e) Geometric model used to estimate the mobile region based on the rotational diffusion (rolling motion) and the contour length of the DNA/RNA duplex. (f) Calculated mobile radius (*R*_mobile_) for the particles with diameters of 100, 500, 1000, and 5000 nm. These geometric parameters are used as inputs to simulations. For the details of the simulation, refer to our previous study (18).

At each RNA site, we assumed an irreversible three-step reaction scheme (Fig. 1b): (1) DNA/RNA hybridization (reaction rate denoted by *k*_on_^DNA/RNA^), (2) RNase H binding to the DNA/RNA duplex (*k*_on_^E^[RNase H]), and (3) RNA hydrolysis followed by dissociation of the hydrolyzed RNA from DNA (*k*_cat_^E^). The values of the reaction rates were set based on our previous study (18) and are listed in Table S1. Reaction events were propagated using the Gillespie algorithm: at each step, an RNA site and a reaction step were chosen with probabilities proportional to the reaction rates, and the simulation time was advanced by an exponentially distributed waiting time determined by the sum of the reaction rates.

To define the region of the RNA surface that can be reached by DNA on the particle, we introduced the accessible radius (*R*_acc_) (Fig. 1c, d). Each ssDNA tether was assumed to access the RNA surface within a distance shorter than its end-to-end length (*L*_0_^DNA^). Using the worm-like chain model, the *L*_0_^DNA^ of a 45-base ssDNA was calculated to be 7.3 nm, corresponding to experimental conditions used in our previous study (18). Combining *L*_0_^DNA^ with the particle radius (*r*_particle_), the accessible radius *R*_acc_ was calculated geometrically from the Pythagorean relation shown in Fig. 1c. Then, the values of *R*_acc_ for the particles with diameters of 100, 500, 1000, and 5000 nm were calculated and plotted in Fig. 1d. The mobile region was defined by the lateral displacement of the particle centroid permitted by a DNA/RNA duplex without exceeding its contour length (Fig. 1e). Following our previous geometric analysis, the total end-to-end distance and the total contour length of the DNA/RNA duplex were set as *L*_0_^DNA/RNA^ = 20.6 nm and *L*_c_ ^DNA/RNA^ = 26.9 nm, respectively, giving a maximum extension of 6.3 nm (18). The corresponding lateral displacement of the particle centroid defines the mobile radius (*R*_mobile_), which specifies the area over which the particle can move while keeping the DNA/RNA duplex within their allowed extension. The values of *R*_mobile_ for the particles with diameters of 100, 500, 1000, and 5000 nm were calculated and plotted in Fig. 1f.

During the simulation, the XY position of the particle was updated after each reaction step based on the distribution of bound RNA sites. For a given set of bound RNA sites, we first calculate the mobile region associated with each bound RNA site (a circle with *R*_mobile_ centered at each RNA site). The allowed positions for the next steps were defined as the overlap of all these mobile regions. If there was an overlapped region, a new particle position was randomly selected within this region, and if there was no overlap, the particle remained at its previous position. The simulation ended when the number of bound RNA sites became zero within the accessible region, corresponding to the detachment from the surface. For each parameter set, 50 independent trajectories were simulated until detachment occurred.

To efficiently simulate long-range motion while keeping the system size manageable, we implemented an adaptive expansion of the RNA surface. The simulation was initialized with a finite square RNA lattice (for example, 600×600 pixels for 100-nm particles; see Table S1 for details). During the simulation, the distance from the position of the particle to the lattice boundary was monitored, and when the position of the particle approached within one quarter of the field size from a given boundary, the lattice was enlarged by adding an extra RNA surface of the same size on that side, generated with the same density and random distribution of RNA sites. The relative coordinates of particle against the newly-added RNA surface were updated to maintain consistency. This adaptive-surface expansion scheme allows long trajectories, including those of large 5000-nm particles, to be computed without allocating an excessively large surface at the start of the simulation.

### Analysis of stepping motion of DNA-nano/microparticle motors

Two-dimensional trajectories of the particle centroid were analyzed using home-built Python code. To mimic experimental localization precision, Gaussian noise with a standard deviation of 2 nm (for both X- and Y-axes) was added to the simulated two-dimensional trajectories before analysis (Fig. 2a). Because the Gillespie algorithm generates events at variable time intervals, the trajectories were first resampled onto time series with constant intervals. We used the constant time intervals of 0.02, 0.02, 0.01, 0.004, 0.004, 0.001, 0.001, 0.0004, 0.0001, and 0.0001 s for *k*_on_^E^[RNase H] = 0.036, 0.144, 0.36, 0.72, 1.44, 3.6, 7.2, 14.4, 36, and 72 s^−1^, respectively. The two-dimensional trajectory data were projected onto X- and Y-axes, and each projection was separately analyzed by using an algorithm established previously (18,24). Briefly, the step-finding algorithm iteratively added steps by calculating the best fit, which minimizes the variance between the raw data and the fit. Then, the variance between the raw data and the counter fit, where step positions were placed between each step position determined by the best fit, was also calculated. For X- and Y-axes, the optimal number of steps and step positions were identified by maximizing the ratio of the sum of squared residuals between the raw data and the counter fit to the sum of squared residuals between the raw data and the best fit. Then, to minimize the under-fitting of the two-dimensional trajectory, we combined the step positions detected only along X-axis (Y-axis) to Y-axis (X-axis). At the same time, to minimize over-fitting (incorrect fitting of a single step as two individual steps) due to the limited detection accuracy of the step positions along X- and Y-axes, the steps detected within ± 5 frames between X- and Y-axes were merged into single steps. These combined step positions were used to re-calculate the fittings for both the X- and Y-projected trajectories, resulting in the final two-dimensional step fitting (Fig. 2b).

**Figure 2.**
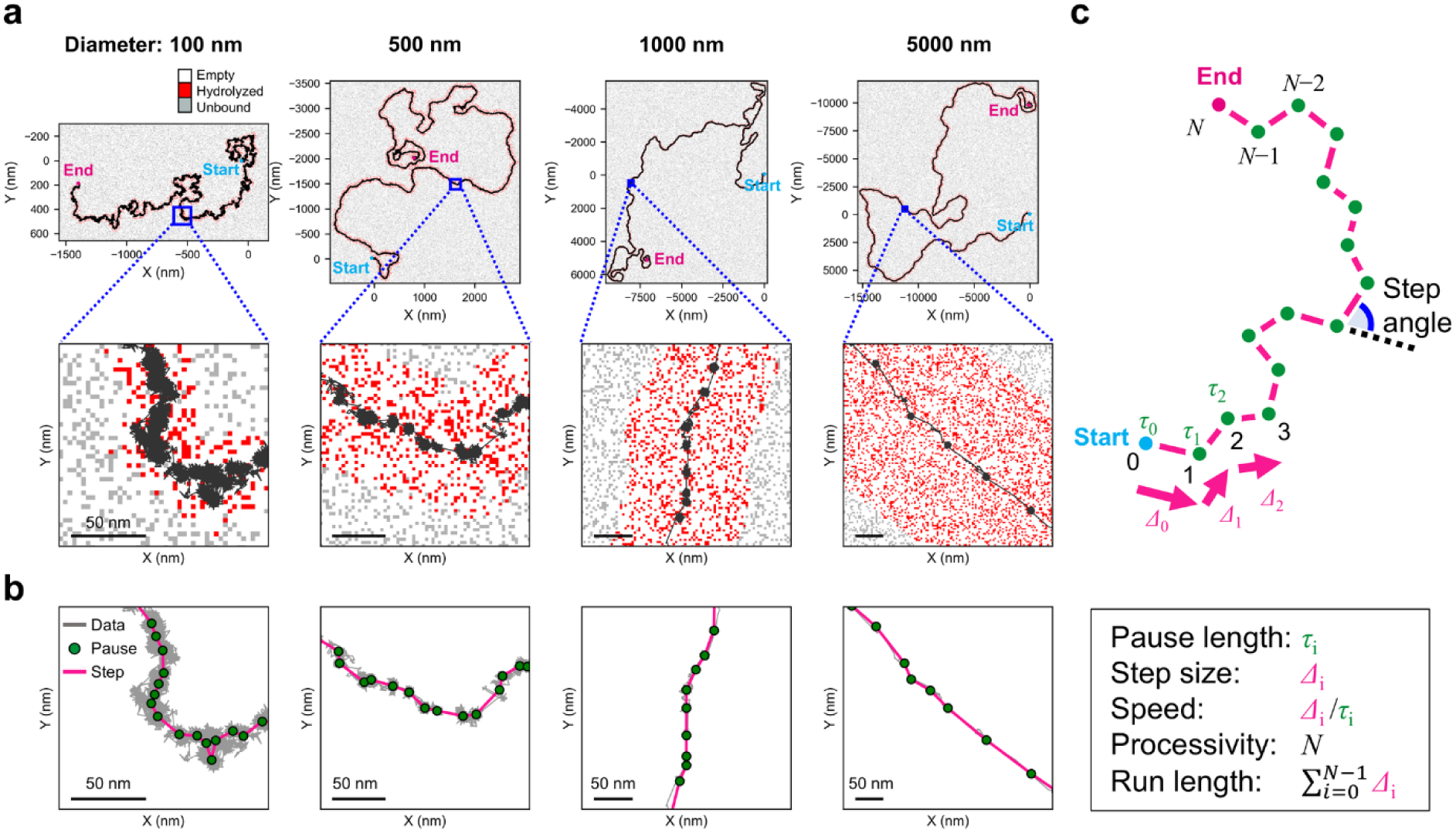
Stepping motions of DNA-nano/microparticle motors observed in simulations and definition of motility and performance metrics. (a) Representative simulated trajectories of the particles with diameters of 100, 500, 1000, and 5000 nm, superimposed on the final spatial distributions of RNA states after the motor run. Red, gray, and white pixels represent hydrolyzed RNA, unbound RNA, and empty (no RNA) sites, respectively. The RNA matrix on the surface was generated using a pixel size normalized by the DNA density on the particle (0.10 molecules nm^−2^). RNAs were randomly distributed according to the RNA density (0.022 molecules nm^−2^). The kinetic parameters were *k*_on_^DNA/RNA^ = 0.8 s^−1^, *k*_on_^E^[RNase H] = 7.2 s^−1^, and *k*_cat_^E^ = 3.0 s^−1^. The top panels show the whole trajectories, and the bottom panels show enlarged views corresponding to the regions indicated by the blue squares in the top panels. Note that Gaussian noise (S.D. = 2 nm for both X- and Y-axes) was added to the simulated trajectories to reproduce the experimental localization precision (18). (b) Examples of pauses (green circles) and steps (magenta lines) detected by the step-finding analysis, superimposed on raw trajectories (gray). (c) Definition of motility and performance metrics used in this study: pause length τ_i_, step size *Δ*_i_, speed *Δ*_i_/*τ*_i_, processivity (total number of steps before detachment) *N*, and run length 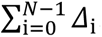. The angle between two consecutive steps was defined as the step angle and used for evaluation of unidirectionality.

### Estimation of motility and performance metrics

Motility and performance metrics—pause length, step size, speed, processivity, run-length, and unidirectionality—were estimated from the two-dimensional step trajectories obtained as described above (Fig. 2b). Pause length (*τ*_i_) was defined as the dwell time between the step *i*−1 and the step *i* (the dwell time between two consecutive steps). Step size (*Δ*_i_) was defined as the spatial displacement associated with step *i*. Speed was defined as *Δ*_i_/*τ*_i_. Unless otherwise noted, the speed in this study refers to the ratio of the step size to the pause length and does not include the time required for stepping (rolling motion). Processivity was defined as the total number of steps (*N*) before detachment. Run-length was defined as the total travel distance along a trajectory before detachment, and was calculated as 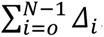. Unidirectionality was quantified based on the distribution of step angle. Each step angle was calculated as the angle between two consecutive steps. For two-dimensional Brownian motion, distribution of the step angle is uniform between −180° and 180°, whereas for perfectly unidirectional ballistic motion, it shows a sharp peak at 0°. Unidirectionality index was defined as (*σ*_uniform_ − *σ*_angle_) / *σ*_uniform_, where *σ*_uniform_ is the standard deviation expected for the uniform distribution (103.9°) and *σ*_angle_ is the standard deviation of the measured step angle distribution. With this definition, the unidirectionality index is 0 when *σ*_angle_ = *σ*_uniform_ (two-dimensional Brownian motion) and 1 when *σ*_angle_ = 0° (perfectly unidirectional ballistic motion).

## Results

### Particle size dependence of speed and run-length reported in the previous experiments

Table 1 summarizes the experimentally determined speed and run-length of DNA-nano/microparticle motors with different materials, DNA/RNA sequences, and particle diameters (16,18). At low [RNase H] between 14 and 720 nM, the speed was nearly proportional to [RNase H], indicating that RNase H binding is the rate limiting. At high [RNase H] between 1440 and 7200 nM, the speed did not depend on [RNase H] and showed the value around 30 nm s⁻¹. Interestingly, the speed at high [RNase H] also did not depend on the particle diameters ranging between 100 nm and 5000 nm, except 96 nm s^−1^ for 100-nm DNA-gold nanoparticle motor with original DNA/RNA sequence at 7200 nM RNase H, which showed apparent two-dimensional Brownian motion caused by imperfect BBR mechanism and a trade-off between the speed and the run-length (see below) (18).

On the other hand, the run-length largely increased as the particle diameter increased. For 5000-nm DNA-silica microparticle motor reported by Yehl et al. (16), the run-length exceeded 200 µm at 1400 nM RNase H, demonstrating highly processive motion. Subsequent experiments using 500-nm DNA-silica nanoparticles showed significantly shorter run-length ranging between 3 and 25 µm, depending on the ionic strength of the buffer solution. In addition, we previously demonstrated that the 100-nm DNA-gold nanoparticle motor showed 2–4 µm run-length, but it decreased to shorter than 1 µm at high [RNase H] for the 100-nm DNA-gold nanoparticle motor with original DNA/RNA sequence (18). Then, we redesigned DNA/RNA sequences to increase DNA/RNA hybridization rate, and successfully improved the trade-off between the speed and the run-length.

### Stepping motions were observed for all particle sizes in simulation

To understand mechanistic origins of the particle size dependence of the speed, run-length, and other performance metrics, we conducted geometry-based kinetic simulations of the DNA-nano/microparticle motors with different diameters of 100, 500, 1000, and 5000 nm (Fig. 1), which cover the range of particle diameters experimentally examined in previous studies (Table 1) (16,18). The detailed algorithm is described in the Materials and Methods. Our simulation captures the interplay between the chemical reaction kinetics and the geometric constraints that collectively determine the motor dynamics. The simulation parameters used in the present study (Table S1) were adopted from our previous study, which successfully reproduced experimentally observed stepping motions of the 100-nm DNA-gold nanoparticle motor (18). Note that unless otherwise noted, we used highest 7200 nM RNase H, which corresponds to the highest RNase H binding rate (*k*_on_^E^[RNase H] = 7.2 s^−1^).

Representative examples of simulated trajectories and underlying RNA states for the motors with different particle diameters are shown in Fig. 2a (corresponding movies of the motion are provided in Supporting Videos S1–4). Note that Gaussian noise (S.D. = 2 nm for both X- and Y-axes) was added to the raw trajectories to mimic the experimental localization precision observed in the previous study (18). Across all particle diameters, the simulated trajectories exhibited clear stepping motions (Fig. 2a, b). In these trajectories, the pauses coincide with frames in which many DNA/RNA duplexes (bound RNA sites) are formed, and the mobile regions of all bound RNA sites do not overlap, so that the particle remains immobilized in place. The steps occur as rapid displacements after the RNA hydrolysis and decrease in the number of bound RNA sites: once a sufficient number of DNA/RNA duplexes are hydrolyzed, the particle becomes mobile within the overlapped mobile region and undergoes a step to a new position, where it becomes immobilized again. These properties became much clear as the particle diameter increased, because the large particle has large number of the RNA sites within the accessible region (Videos S1–4).

### Particle size dependence of steps size, pause length, and speed

To characterize the stepping motion quantitatively, we next applied a step-finding algorithm to the simulated two-dimensional trajectories (Materials and Methods, Fig. 2b) (18,24). The fitted trajectories allowed us to define a set of motility and performance metrics (Fig. 2c): the pause length *τ*_i_, step size *Δ*_i_, speed *Δ*_i_/*τ*_i_, processivity *N* (total number of steps before detachment), and run-length 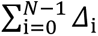. In addition, the angle between two consecutive steps was defined as step angle, and unidirectionality was quantified from the distribution of step angle (see Materials and Methods for details). It should be noted that the speed defined above does not include the duration required for stepping (rolling motion) driven by the rotational diffusion of the particle. Thus, the speed can be overestimated, especially when the stepping duration becomes comparable to or longer than the pause length. The effect of rotational diffusion is discussed later, where the stepping duration is incorporated into the speed evaluation.

By analyzing simulated trajectories, we obtained the motility and performance metrics for all combinations of the particle diameters (100, 500, 1000, and 5000 nm) and DNA/RNA hybridization rates (*k*_on_^DNA/RNA^) used in the previous experiments (Table 2) (16,18). We first considered the DNA/RNA hybridization rate corresponding to the original sequence (*k*_on_^DNA/RNA^ = 0.2 s^−1^, Fig. 3a). As the particle diameter increased from 100 nm to 5000 nm, the step size increased (Fig. 3a, top), consistent with the particle diameter dependences of *R*_acc_ and *R*_mobile_ derived from the geometric model (Fig. 1d, f). In addition, the pause length also increased as the particle diameter increased, indicating that large particles tend to remain immobilized for long periods between the steps (Fig. 3a, middle). This trend arises because the accessible region of a large particle contains a large number of RNA sites on the surface, leading to the formation of a large number of DNA/RNA duplexes. Consequently, more DNAs and RNAs should undergo the hybridization-hydrolysis cycle before mechanical constraints are relieved to allow next step, resulting in the long pauses. We also analyzed the simulation data using the DNA/RNA hybridization rate corresponding to the newly-designed sequence (*k*_on_^DNA/RNA^ = 0.8 s^−1^, Fig. 3b) (18). For all particle diameters, the pause length and the step size showed similar diameter dependence with those using the original sequence.

**Figure 3.**
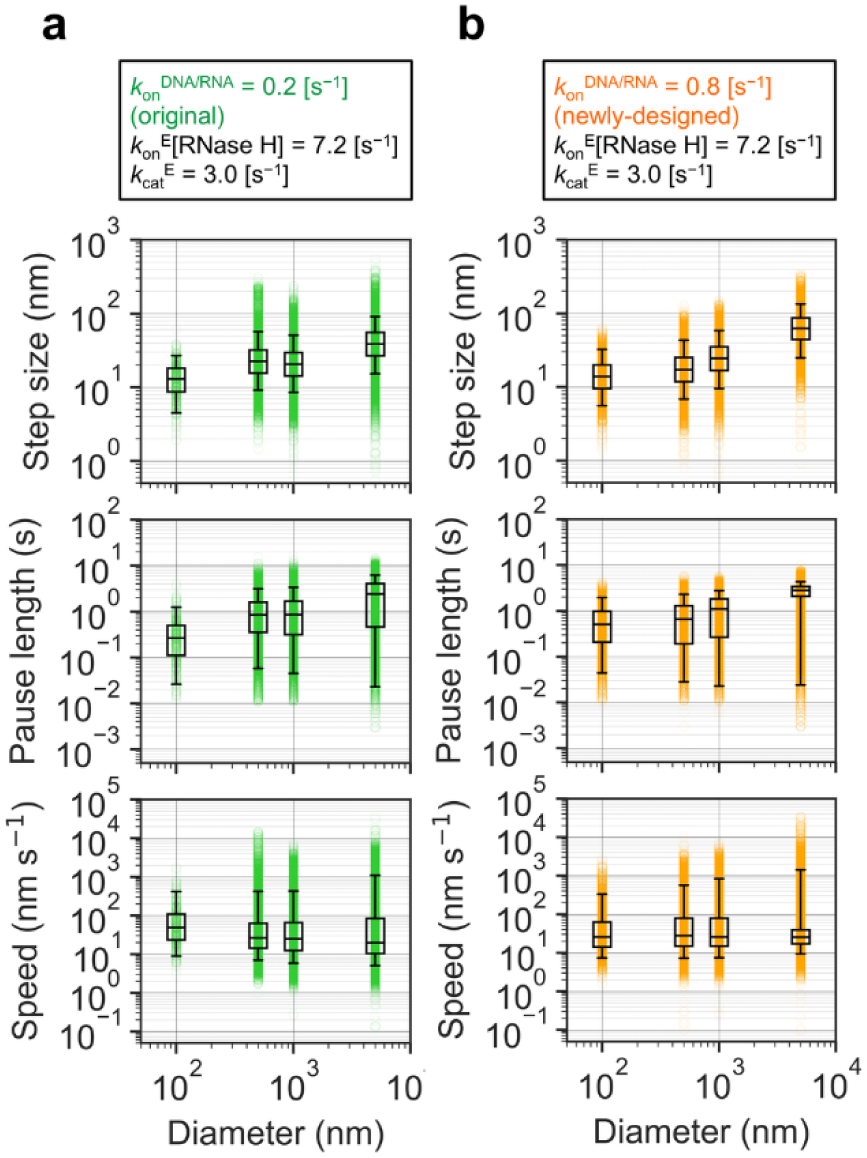
Particle size dependence of step size, pause length, and speed. (a) Pause length (top), step size (middle), and speed (bottom) obtained from simulations of the particles with diameters of 100, 500, 1000, and 5000 nm using *k*_on_^DNA/RNA^ = 0.2 s^−1^, *k*_on_^E^[RNase H] = 7.2 s^−1^, and *k*_cat_^E^ = 3.0 s^−1^. *k*_on_^DNA/RNA^ = 0.2 s^−1^ corresponds to the original DNA/RNA sequence used in the previous study (18). (b) Results using *k*_on_^DNA/RNA^ = 0.8 s^−1^, corresponding to the newly-designed DNA/RNA sequence reported in the previous study (18). In all panels, box plots show the interquartile range (IQR), the horizontal lines indicate the medians, and whiskers represent the 5th–95th percentiles. Individual data points from 50 simulated trajectories are shown as semi-transparent circles.

**Table 2.** Motility and performance metrics of DNA-nano/microparticle motors with different particle sizes and *k*_on_^DNA/RNA^, estimated by geometry-based kinetic simulations^a^.

| Particle diameter (nm) | $k_{\text{on}}^{\text{DNA/RNA}}$ ($\text{s}^{-1}$) | Step size (nm) | Pause length (s) | Speed ($\text{nm s}^{-1}$) | Processivity | Run-length ($\mu\text{m}$) | Unidirectionality |
| --- | --- | --- | --- | --- | --- | --- | --- |
| 100 | 0.2<sup>b</sup> | 12.9 | 0.26 | 48.1 | 14 | 0.21 | 0.11 |
| 500 |  | 22.3 | 0.84 | 26.4 | 767 | 21 | 0.33 |
| 1000 |  | 20.5 | 0.85 | 25.0 | 1782 | 43 | 0.46 |
| 5000 |  | 38.6 | 2.37 | 19.6 | 1619 | 72 | 0.65 |
| 100 | 0.8<sup>c</sup> | 13.8 | 0.51 | 25.8 | 124 | 2.0 | 0.29 |
| 500 |  | 17.1 | 0.66 | 27.8 | 653 | 13 | 0.58 |
| 1000 |  | 24.3 | 1.10 | 25.9 | 800 | 22 | 0.65 |
| 5000 |  | 62.5 | 2.78 | 25.4 | 732 | 51 | 0.75 |

<sup>a</sup> $[\text{RNase H}] = 7200 \text{ nM}$ , $k_{\text{on}}^{\text{E}}[\text{RNase H}] = 7.2 \text{ s}^{-1}$ , and $k_{\text{cat}}^{\text{E}} = 3.0 \text{ s}^{-1}$
<sup>b</sup> Corresponds to original DNA/RNA sequence in the experiment
<sup>c</sup> Corresponds to newly-designed DNA/RNA sequence in the experiment

Interestingly, because of the trade-off between the step size and the pause length, the median speed did not change largely for *k*_on_^DNA/RNA^ = 0.2 s^−1^ (original DNA/RNA sequence, Table 2, Fig. 3a, bottom). This trade-off became more clear for *k*_on_^DNA/RNA^ = 0.8 s^−1^ (newly-designed DNA/RNA sequence), and resulted in nearly constant speed in the range of 25∼28 nm s^−1^ (Table 2, Fig. 3b, bottom). As results, our simulation quantitatively reproduced particle size-independent speed (∼30 nm s^−1^) observed in the experiments (Table 1).

### Particle size dependence of processivity and run-length correlates with detachment modes

We next asked why the processivity and run-length exhibits strong dependence on the particle diameter (Table 1 and 2). In our previous experiments of the 100-nm DNA-gold nanoparticle motor (18), we showed that the processivity and run-length strongly depended on the balance among the DNA/RNA hybridization rate (*k*_on_^DNA/RNA^), the RNase H binding rate (*k*_on_^E^[RNase H]), and the RNA hydrolysis rate (*k*_cat_^E^). When *k*_on_^DNA/RNA^ is much larger than *k*_on_^E^[RNase H] and *k*_cat_^E^, the processivity and run-length approach an upper limit caused by an entrap detachment, in which the motor is entrapped within the region where the surrounding RNAs are already hydrolyzed and eventually forced to detach from the surface (Fig. 4a, c). Note that in the case of the entrap detachment, upper limit of the processivity and run-length depends on the unidirectionality of the motion (see below). In contrast, when *k*_on_^DNA/RNA^ is much smaller than *k*_on_^E^[RNase H] and *k*_cat_^E^, the processivity and run-length are decreased by a stochastic detachment, which occurs when all DNA/RNA duplexes are hydrolyzed stochastically before the formation of new DNA/RNA duplexes (Fig. 4b, d).

**Figure 4.**
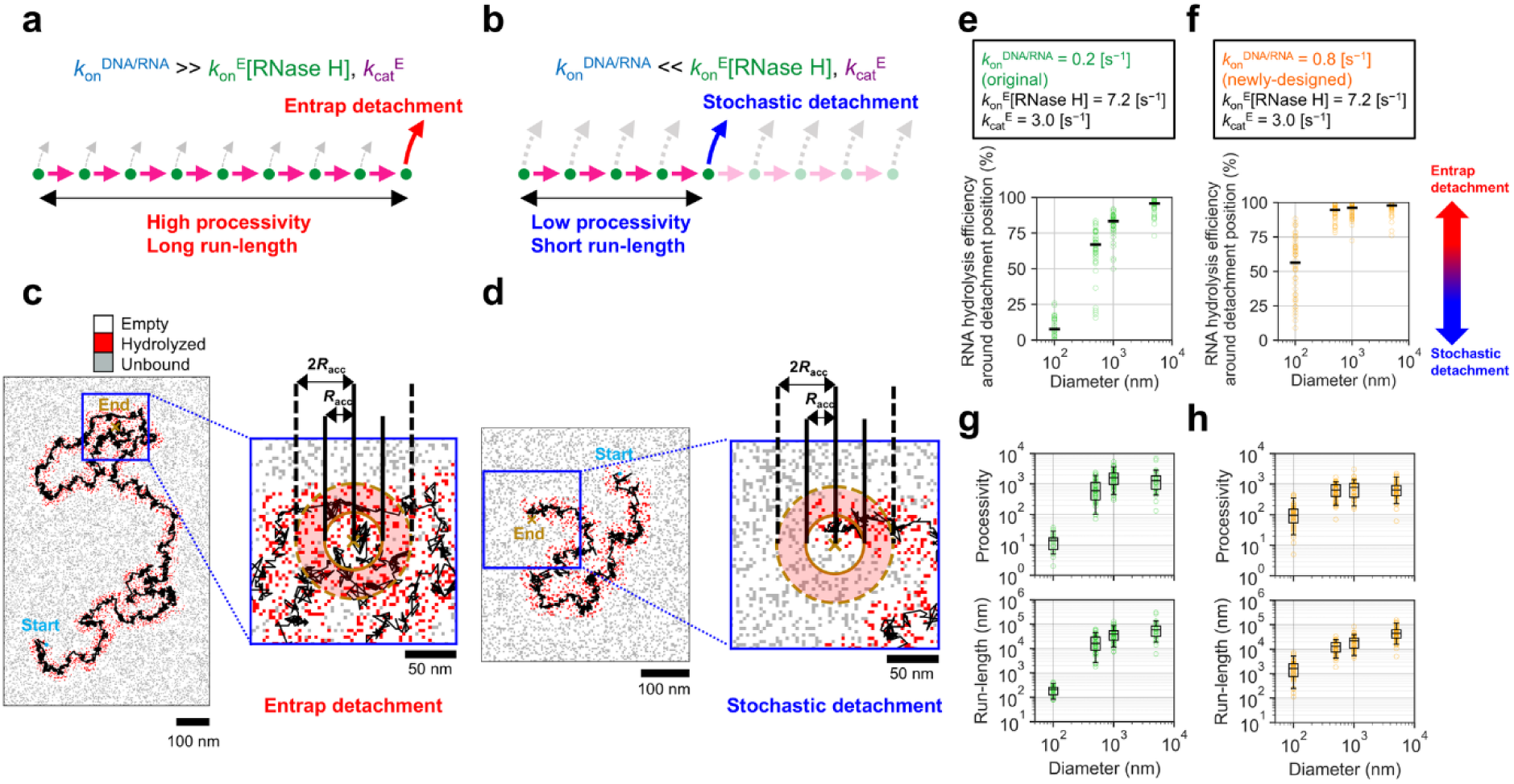
Entrap and stochastic detachments and particle size dependence of RNA hydrolysis efficiency around detachment position, processivity, and run-length. (a, b) Schematic illustrations of two different mechanisms of detachment observed in this study, entrap detachment (a) and stochastic detachment (b). (c, d) Representative examples of RNA state distributions around detachment positions for entrap detachment (c) and stochastic detachment (d). Red, gray, and white pixels represent hydrolyzed RNA, unbound RNA, and empty (no RNA) sites, respectively. The kinetic parameters are *k*_on_^DNA/RNA^ = 0.8 s^−1^, *k*_on_^E^[RNase H] = 7.2 s^−1^, and *k*_cat_^E^ = 3.0 s^−1^. Red shaded donut-shaped regions indicate the areas between *R*_acc_ and 2*R*_acc_ from the detachment positions. (e, f) RNA hydrolysis efficiency around the detachment positions for particles with diameters of 100, 500, 1000, and 5000 nm, obtained using *k*_on_^DNA/RNA^ = 0.2 s^−1^ (e) or 0.8 s^−1^ (f). Black solid horizontal lines show the median values. High and low values correspond to entrap and stochastic detachments, respectively. (g, h) Particle diameter dependence of the processivity (top) and run-length (bottom) under the same kinetic conditions as in (e) and (f), respectively. Box plots show the interquartile range (IQR), the horizontal lines indicate the medians, and whiskers represent the 5th–95th percentiles. Individual data points from 50 simulated trajectories are shown as semi-transparent circles.

To elucidate the origin of the particle diameter dependence of the processivity and run-length (Table 1 and 2), we analyzed how frequently the motors detached via two distinct detachment modes (entrap and stochastic) under different particle diameters and *k*_on_^DNA/RNA^. To analyze two detachment modes in a quantitative manner, we estimated the RNA hydrolysis efficiency around the detachment position of each trajectory by calculating the fraction of hydrolyzed RNA within the donut-shaped region between *R*_acc_ and 2*R*_acc_ (Fig. 4c, d). High and low RNA hydrolysis efficiencies correspond to the entrap and stochastic detachments, respectively. For 100-nm particle, increasing *k*_on_^DNA/RNA^ from 0.2 to 0.8 s^−1^ markedly increased the fraction of the entrap detachment and increased the processivity and run-length nearly 10 times (Fig. 4e-h, Table 2), reproducing our previous experiments (Table 1) (18).

Interestingly, the present simulation further revealed that the detachment mode also highly depends on the particle diameter. Even with the low DNA/RNA hybridization rate (*k*_on_^DNA/RNA^ = 0.2 s^−1^), the RNA hydrolysis efficiency around the detachment position largely increased as particle diameter increased (Fig. 4e), indicating that large particle exhibits a high fraction of the entrap detachment. This change in the detachment modes explains the particle diameter dependence of the processivity and run-length (Fig. 4g, h). For 100-nm particle with *k*_on_^DNA/RNA^ = 0.2 s^−1^, stochastic detachment was dominant and resulted in low processivity and short run-length (Fig. 4g). As the particle diameter increased, stochastic detachment became minor, and the processivity rapidly increased and saturated around 1000 steps. On the other hand, because the step size gradually increased with the particle diameter (Table 2, Fig. 3), the run-length gradually increased accordingly. When *k*_on_^DNA/RNA^ was increased to 0.8 s^−1^, the ratio of entrap detachment increased also for 100-nm particle, resulting in a 10-fold increase in processivity and run-length (Fig. 4h), compared with those at *k*_on_^DNA/RNA^ = 0.2 s^−1^ (Fig. 4g). In contrast, the processivity and run-length of 500, 1000, 5000-nm particles were less sensitive to the change in DNA/RNA hybridization rate (Fig. 4g, h, Table 2), consistent with the result that the entrap detachment is already major for these particle diameters (Fig. 4e, f).

### Particle size dependence of unidirectionality correlates with number of bound RNA sites

We next investigated how unidirectionality of the stepping motion depends on the particle diameter (Fig. 5). The step angle, defined as the angle between two consecutive steps, showed characteristic changes in its distribution as the particle diameter increased (Fig. 5a). For all diameters, the step angle distributions were biased toward the forward direction, indicating that the motors operate as the BBR. The polar plots gradually changed from relatively-broad lobes for 100-nm particles to sharply focused forward peaks for 5000-nm particles. This tendency was captured quantitatively by the unidirectionality index defined using the normalized standard deviation of the step angle distribution (Materials and Methods), demonstrating a clear diameter dependence (Fig. 5b, Table 2).

**Figure 5.**
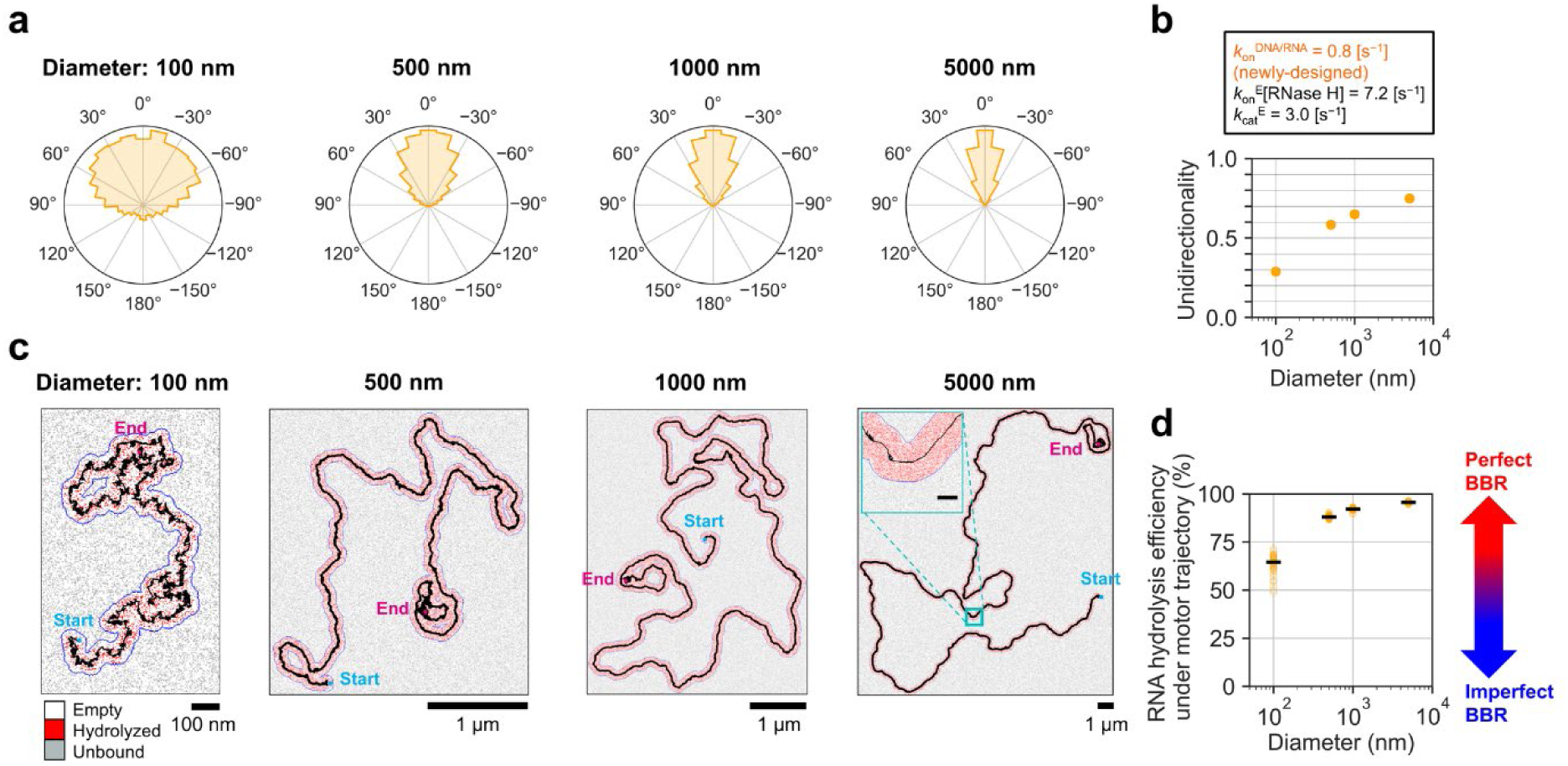
Particle size dependence of step angle distribution, unidirectionality, and RNA hydrolysis efficiency under motor trajectory. (a) Polar plots of step angles for particles with diameters of 100, 500, 1000, and 5000 nm. The forward direction is defined as 0°. The kinetic parameters are *k*_on_^DNA/RNA^ = 0.8 s^−1^, *k*_on_^E^[RNase H] = 7.2 s^−1^, and *k*_cat_^E^ = 3.0 s^−1^. (b) Particle diameter dependence of unidirectionality index, calculated from the standard deviation of step angle distribution normalized by the standard deviation of a uniform angular distribution. (c) Representative trajectories for each particle diameter, superimposed on the final spatial distribution of RNA states after the motor run. Red, gray, and white pixels represent hydrolyzed RNA, unbound RNA, and empty (no RNA) sites, respectively. Blue contours indicate the regions under the motor trajectory. (d) Particle diameter dependence of RNA hydrolysis efficiency under the motor trajectory. High and low values correspond to perfect and imperfect BBR, respectively.

We then analyzed RNA hydrolysis efficiency under the motor trajectory (Fig. 5c, d), which serves as a quantitative measure of the perfectness of the BBR mechanism. The efficiency increased with the particle diameter and showed a positive correlation with the unidirectionality index (Fig. 5b), indicating that high RNA hydrolysis efficiency under the motor trajectory is associated with highly biased forward motion. This result is consistent with the BBR mechanism: as RNA sites under the motor are more completely hydrolyzed, the probability of returning to positions previously visited decreases, enhancing directional bias. We also found that the perfectness of the BBR mechanism alone cannot fully account for the high unidirectionality observed for large particles. In our previous study of 100-nm DNA-gold nanoparticle motor, we showed that the maximum values of the unidirectionality index were ∼0.4 for experiments and ∼0.5 for simulations, even under conditions of low [RNase H] where the RNA hydrolysis efficiency under the motor trajectory was more than 90% (18). These values are much lower than the unidirectionality index (0.75) obtained for 5000-nm particle in the present simulation (with *k*_on_^DNA/RNA^ = 0.8 s^-1^, Fig. 5b, Table 2).

To uncover the origin of particle diameter dependence of the unidirectionality, we next analyzed how the spatial distribution of bound RNA sites just before step correlates with the step direction (Fig. 6). In our model, the possible positions of the particle after a step are restricted to the overlap of the mobile regions of all bound RNA sites (Fig. 6a, green square pixels). As an effective descriptor of this constraint, we focused on the centroid of all bound RNA sites within the accessible region at the frame just before stepping (Fig. 6a, green crosses). Based on this definition, we quantified three angular quantities: step angle (angle between previous and current step directions, Δ*θ*_prev,current_, Fig. 6b, top left), angle between previous step direction and direction from particle centroid toward centroid of all bound RNA sites (Δ*θ*_prev,bound_, Fig. 6b, middle left), and angle between direction from particle centroid toward centroid of all bound RNA sites and current step direction (Δ*θ*_bound, current_, Fig. 6b, bottom left).

**Figure 6.**
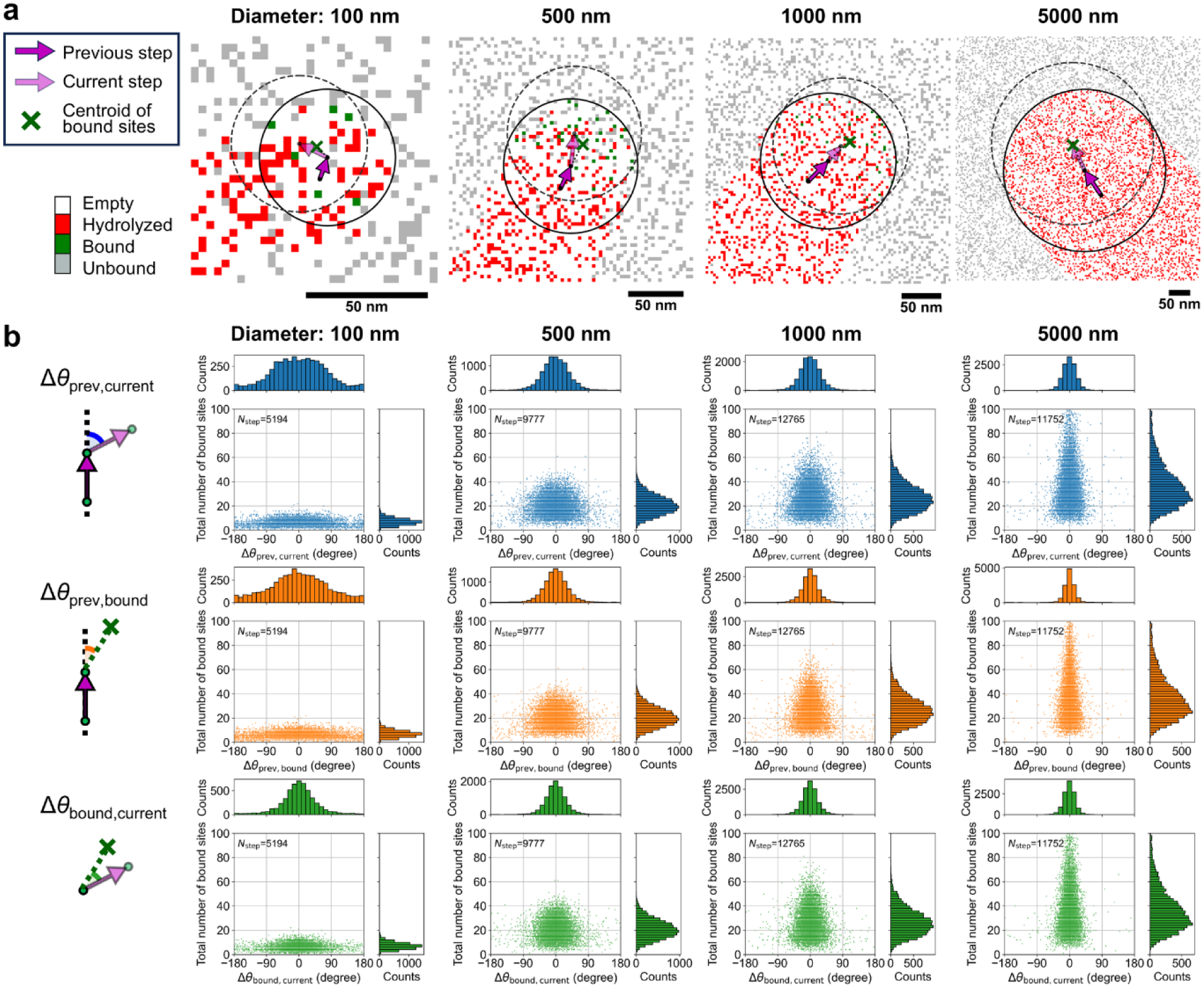
Particle size dependence of spatial distribution of bound RNA sites within accessible region and correlation between total number of bound RNA sites and step direction. (a) Representative RNA state distributions at the frame just before the step for particles with diameters of 100, 500, 1000, and 5000 nm. Red, green, gray, and white pixels indicate hydrolyzed RNA, bound RNA, unbound RNA, and empty (no RNA) sites, respectively. Black solid and dashed circles denote the accessible regions just before and after the steps, respectively. Purple and transparent-purple arrows indicate directions of the previous and current steps, respectively. Green crosses indicate the centroid of all bound RNA sites just before the steps. (b) Correlation between total number of bound RNA sites within accessible region and Δ*θ*_prev,current_ (angle between previous and current step directions, i.e., step angle), Δ*θ*_prev,bound_ (angle between previous step direction and direction from particle centroid toward centroid of all bound RNA sites), or Δ*θ*_bound,current_ (angle between direction from particle centroid toward centroid of all bound RNA sites and current step direction). Data were obtained from 50 simulated trajectories for each particle diameter. The kinetic parameters are *k*_on_^DNA/RNA^ = 0.8 s^−1^, *k*_on_^E^[RNase H] = 7.2 s^−1^, and *k*_cat_^E^ = 3.0 s^−1^.

First, we investigated correlation between Δ*θ*_prev,current_ and the total number of bound RNA sites for different particle diameters (Fig. 6b, top). For the 100-nm particle, total number of bound RNA sites was low and Δ*θ*_prev,current_ showed broad distribution, although biased to forward (between ±90°). As the particle diameter increased, the fraction of large total number of bound RNA sites increased, and importantly, Δ*θ*_prev,current_ showed narrow distribution and a sharp peak around 0°. Considering the RNA density used in the simulation (Table S1), average number of RNA sites within the accessible region is 54, 256, 508, and 2524 for particles with diameters of 100, 500, 1000, and 5000 nm, respectively. Therefore, as the particle diameter increases, the upper limit of total number of bound RNA sites increases. Evidently, large total number of bound RNA sites strongly correlates with high unidirectionality.

Next, correlations between Δ*θ*_prev,bound_ and total number of bound RNA sites were analyzed (Fig. 6b, middle, bottom). Δ*θ*_prev,bound_ showed almost identical correlation with that of Δ*θ*_prev,current_. For large particles, distributions of Δ*θ*_prev,bound_ showed narrow peaks around 0°, indicating that the centroid of all bound RNA sites just before the step tends to lie ahead of the particle along the direction of previous step. This sharp forward bias originates from the geometry of the accessible region. The difference between the circular accessible regions before a step (Fig. 6a, black solid circles) and the region after the step (Fig. 6a, black dashed circles) defines a crescent-shaped area biased along the previous step direction. As the number of bound RNA sites increases, the direction of the centroid of all bound RNA sites tends to converge toward the previous step direction. In contrast, for the 100-nm particle, the distribution was broad, reflecting small number of bound RNA sites and weak forward averaging.

On the other hand, Δ*θ*_bound,current_ showed slightly different correlation with that of Δ*θ*_prev,current_. Especially, for 100-nm particle, distribution of Δ*θ*_bound,current_ showed narrower peak around 0° than that of Δ*θ*_prev,current_ (Fig. 6b, bottom left). This indicates a robust correlation between the centroid direction and the current step direction. This result confirms that the current step is taken toward the region where bound RNA sites are concentrated. Taken together, these results indicate that high unidirectionality of large particles arises not only from almost complete RNA hydrolysis under the motor trajectory but also from an averaging effect of the large number of bound RNA sites on the step directions.

### Improvements in all kinetic parameters are required to achieve speeds exceeding 100 nm s⁻¹ while maintaining other performance metrics

The detailed analyses described above showed that the speed (∼30 nm s^−1^) is essentially independent of the particle diameter, whereas the run-length exhibits a pronounced diameter dependence (Figs. 3–6, Table 2). To identify the kinetic parameters primarily controlling the speed, and the requirements to achieve the speed larger than 100 nm s^−1^, we systematically varied the three reaction rates, *k*_on_^DNA/RNA^, *k*_on_^E^[RNase H], and *k*_cat_^E^, focusing on the 100-nm particle. For each kinetic parameter set, we estimated the speed and the run-length from the stepping trajectories and plotted them in the speed–run-length planes (Fig. 7).

**Figure 7.**
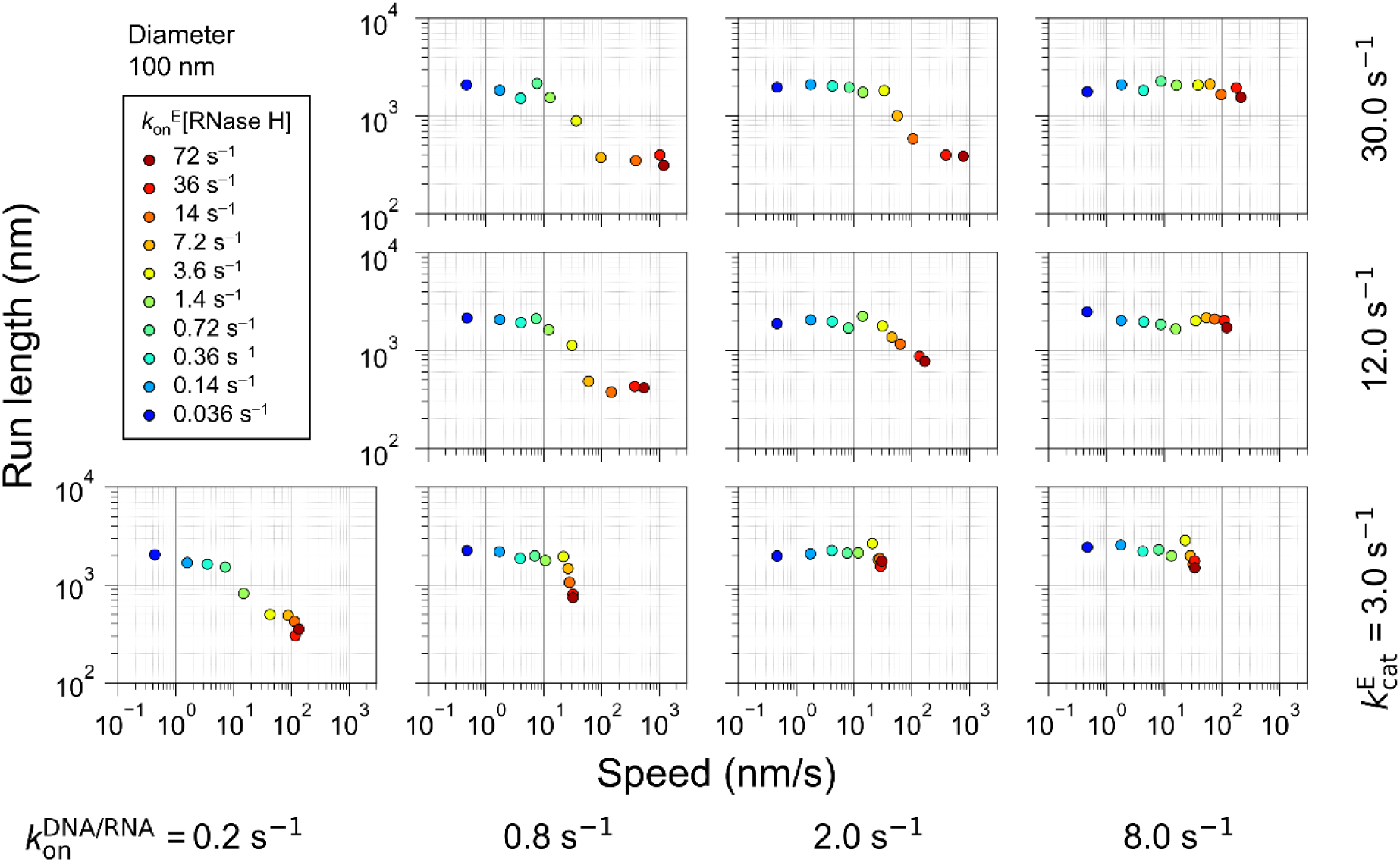
Relationship between speed and run-length under various kinetic parameters for the motor with 100 nm particle. Scatter plots of median speed versus mean run-length obtained from 50 simulations of the motor with 100 nm particle under various kinetic conditions. Individual data points correspond to different combinations of DNA/RNA hybridization rates (*k*_on_^DNA/RNA^, from left to right panels), RNase H binding rates (*k*_on_^E^[RNase H], symbols with different colors in each panel), and RNA hydrolysis rates (*k*_cat_^E^, from bottom to top panels).

Under the kinetic conditions corresponding to the original and newly-designed DNA/RNA sequences used in the previous sections (*k*_on_^DNA/RNA^ = 0.2 and 0.8 s^−1^, respectively, and *k*_cat_^E^ = 3.0 s^−1^), we observed a clear trade-off between the speed and the run-length as *k*_on_^E^[RNase H] increased (Fig. 7, bottom left). Increase in *k*_on_^E^[RNase H] accelerated RNase H binding, shortened the pause length, and increased the speed. However, it resulted in short run-length because the stochastic detachment became dominant (Fig. 4). Simultaneous increases in *k*_on_^E^[RNase H] and *k*_cat_^E^ also resulted in similar trade-off (Fig. 7, left columns): fast RNA hydrolysis further shortened the pause length and increased the speed, but decreased the run-length due to the stochastic detachment.

On the other hand, when *k*_on_^DNA/RNA^, *k*_on_^E^[RNase H], and *k*_cat_^E^ were increased simultaneously, the trade-off between the speed and the run-length was improved (Fig. 7, top right). With large *k*_on_^DNA/RNA^, DNA/RNA duplexes are rapidly formed, which suppresses stochastic detachment even under the condition of large *k*_on_^E^[RNase H] (fast RNase H binding) and large *k*_cat_^E^ (fast RNA hydrolysis). Consequently, the data points shift toward the upper-right region of the speed–run-length plane, where both high speed and long run-length are achieved. Notably, at *k*_on_^DNA/RNA^ = 8.0 s^−1^, *k*_on_^E^[RNase H] = 72 s^−1^, and *k*_cat_^E^ = 30.0 s^−1^, the 100-nm particle motor showed the speed of ∼200 nm s^−1^ and the run-length of ∼2 μm (Fig. 7, top right), both comparable to those of a motor protein kinesin-1 (25–27).

In addition, plots of speed versus processivity and speed versus unidirectionality index for the same kinetic parameter sets exhibited similar trade-off relationships (Figs. S1 and S2, respectively): the conditions that yielded high speed generally showed low processivity and low unidirectionality, particularly with small *k*_on_^DNA/RNA^. To achieve high speed and high processivity or high unidirectionality simultaneously, all three kinetic parameters need to be increased simultaneously.

Interestingly, maximum values of the run-length (∼2 μm), the processivity (∼1000), and the unidirectionality index (∼0.5) were not improved even in the case of large values of all three kinetic parameters (Fig. 7, Figs. S1 and S2). These results indicate that whereas the maximum speed is primarily governed by the kinetic parameters, the maximum values of run-length and processivity are limited by the entrap detachment (Fig. 4) caused by a fundamental limitation of the two-dimensional BBR motor, the imperfectness of the unidirectionality (see Discussion below).

### Diffusion-limited stepping restricts maximum speed of micron-sized particle motors

The kinetic parameter sweep shown in Fig. 7 indicated that, for 100-nm particle, simultaneous acceleration of DNA/RNA hybridization, RNase H binding and RNA hydrolysis can yield high speeds of ∼200 nm s^−1^ while maintaining long run-length of ∼2 μm. In the simulations described above, we assumed that steps occur instantaneously and that the speed is solely determined by the ratio of step size (*Δ*_i_) to pause length (*τ*_i_) (Fig. 2). However, in reality, the step itself requires a finite time because the particle must undergo rotational diffusion (rolling motion, Fig. 8a). Therefore, we next asked whether the duration required for rolling motion (stepping duration, *τ*_rot_) can become rate-limiting of the motor motion.

**Figure 8.**
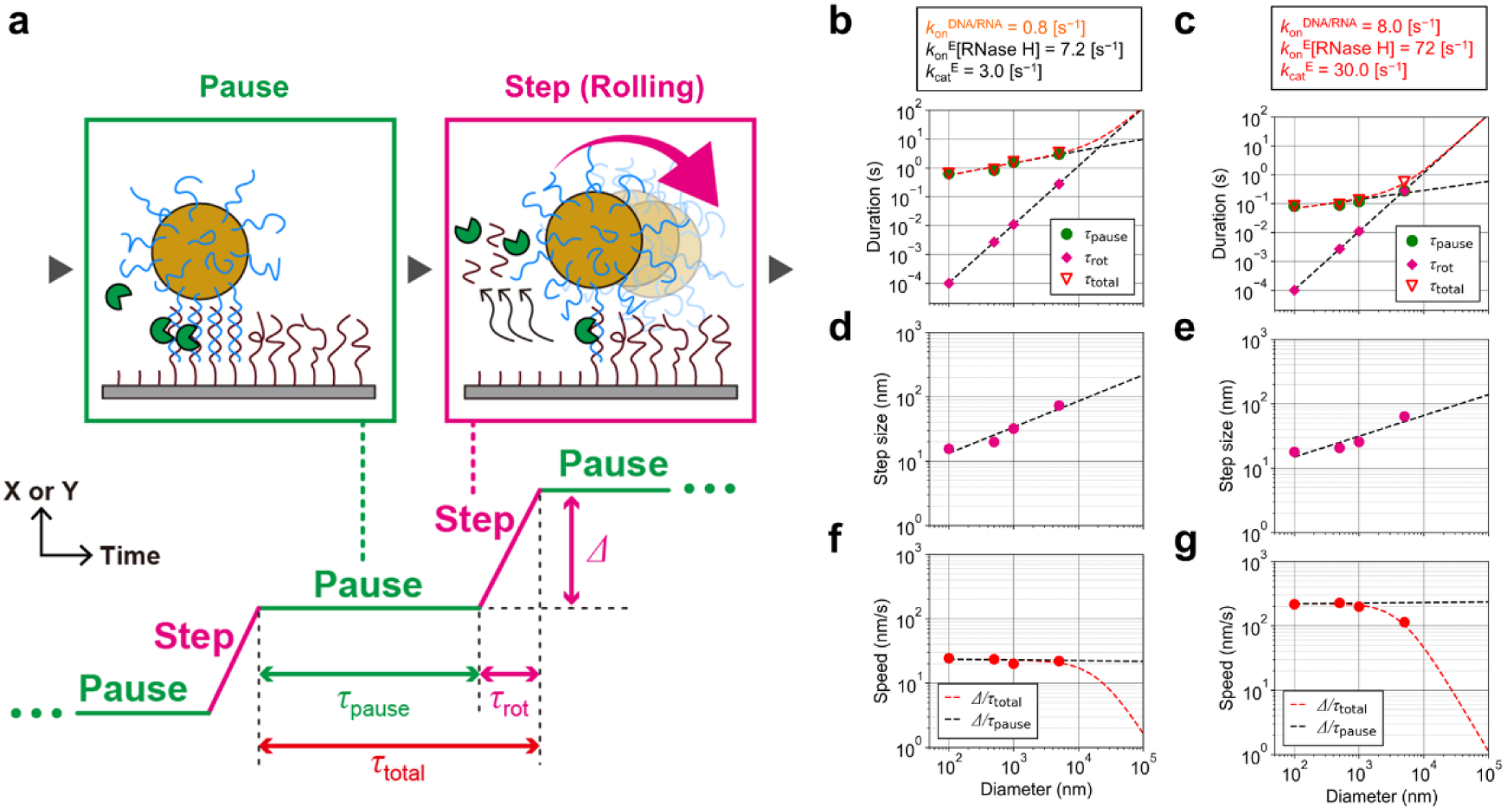
Effect of particle-size dependent rotational diffusion on the speed under different kinetic conditions. (a) Schematic illustration of median pause length (*τ*_pause_), stepping duration (*τ*_rot_) estimated from the first passage time of rotational diffusion, and median step size (*Δ*). (b, c) Particle diameter dependence of *τ*_pause_ (green circles), *τ*_rot_ (pink diamonds), and total cycle time *τ*_total_ (*τ*_pause_ + *τ*_rot_) (red inverted triangles) under two kinetic conditions: (b) *k*_on_^DNA/RNA^ = 0.8 s^−1^, *k*_on_^E^[RNase H] = 7.2 s^−1^, and *k*_cat_^E^ = 3.0 s^−1^, (c) *k*_on_^DNA/RNA^ = 8.0 s^−1^, *k*_on_^E^[RNase H] = 72 s^−1^, and *k*_cat_^E^ = 30 s^−1^. Black dashed lines indicate linear fits of *τ*_pause_ or *τ*_rot_ in the log–log plots. Red dashed lines are sums of the two black dashed lines. (d, e) Particle diameter dependence of step size (*Δ*) under the same kinetic conditions as in (b) and (c), respectively. Purple circles indicate median values obtained from the simulations, and black dashed lines indicate linear fits in the log–log plots. (f, g) Speed calculated from the step size (b) and the duration (a). Red circles and red dashed lines correspond to *Δ*/*τ*_total_ and black dashed lines correspond to *Δ*/*τ*_pause_ under the same kinetic conditions as in (b) and (c), respectively.

To address this question, we estimated *τ*_rot_ as the first-passage time of the rotational diffusion required for the particle to move a distance corresponding to the mobile radius (Fig. 1e, f), based on the rotational diffusion coefficient (18). We then compared *τ*_rot_ with the median pause length (*τ*_pause_) obtained from the geometry-based kinetic simulation for each particle diameter (Fig. 3 and Fig. 8b, c). As a result, under the kinetic condition used in most of this study (*k*_on_^DNA/RNA^ = 0.8 s^−1^, *k*_on_^E^[RNase H] = 7.2 s^−1^, *k*_cat_^E^ = 3.0 s^−1^, Fig. 8b), *τ*_rot_ increased nonlinearly with the particle diameter but remained much shorter than *τ*_pause_ over the entire range from 100 to 5000 nm. Consequently, effective speed calculated by the ratio of median step size (*Δ*, Fig. 3 and Fig. 8d) to total cycle time (*τ*_total =_ *τ*_pause_ + *τ*_rot_) was almost indistinguishable from the speed simply estimated as *Δ*/*τ*_pause_ (Fig. 8f). This result indicates that rotational diffusion does not limit the speed under these conditions, even for micron-sized particle motors.

In contrast, for the condition that yields a very high speed of ∼200 nm s^-1^ (*k*_on_^DNA/RNA^ = 8.0 s^−1^, *k*_on_^E^[RNase H] = 72 s^−1^, and *k*_cat_^E^ = 30.0 s^−1^; Fig. 7), the situation changed (Fig. 8c, e, g). Under this condition, *τ*_pause_ shortened 10 times but *τ*_rot_ remained the same. For the 100-nm particle, *τ*_rot_ was still much smaller than *τ*_pause_, and the speed remained close to the predicted value of ∼200 nm s^−1^. However, for the 5000-nm particle, *τ*_rot_ became comparable to *τ*_pause_, indicating that the duration for the rotational diffusion and that for the chemical reactions contributed similarly to the total cycle time. As a result, the speed was reduced to roughly half, ∼100 nm s^−1^ (Fig. 8g).

## Discussion

### Nano-sized DNA-particle motors are essential to rival motor proteins

Our simulations clarify why the speed (∼30 nm s^−1^) of DNA-nano/microparticle motors remains independent of the particle diameter (Figs. 3 and 7). This speed is comparable to that of a BBR linear motor protein chitinase which moves along single chitin chains at ∼50 nm s^−1^, driven by chitin degradation from the end (14,28). On the other hand, typical ATP-driven linear motor proteins are substantially faster than the chitinase. For example, kinesin-1 moves at the speed of 600–800 nm s^−1^ at saturating ATP concentration (12,25), and myosin V shows a speed of ∼1 µm s^−1^ *in vitro* and in living cells (13,29). In these motor proteins, elementary processes of ATP hydrolysis reaction cycle (ATP binding, cleavage of phosphate bond, ADP and phosphate releases) which cause the pauses proceeds on the order of 1–10 ms (25), whereas the pause lengths of the DNA-nano/microparticle motors are still longer than 100 ms (Figs. 3 and 8) (18). This difference highlights that further acceleration of the chemical reaction cycle (or shortening of the pause length) is required to fill the speed gap.

By systematically varying kinetic parameters, we found that 100-nm DNA-nanoparticle motor can reach a speed of ∼200 nm s^−1^ while maintaining micrometer-scale run-length if DNA/RNA hybridization, RNase H binding, and RNA hydrolysis are accelerated simultaneously (Fig. 7). Experimentally, the DNA/RNA hybridization rate (*k*_on_^DNA/RNA^) can be elevated by redesigning DNA/RNA sequences. Our previous work with repetitive and GC-rich sequences showed that sequence redesign substantially accelerated DNA/RNA hybridization and improved the trade-off between the speed and the run-length (18). Our current results indicate that further optimization in this direction is a promising route to further improving speed. The RNase H binding rate (*k*_on_^E^[RNase H]) can be easily increased by increasing [RNase H]. In contrast, the catalytic rate (*k*_cat_^E^) highly depends on the properties of the enzymes used. While RNase H is the first choice for DNA/RNA-based BBR motors (16), recent demonstrations of a peptide-based BBR motors driven by a protease (19) and an RNA-free, DNA-based BBR motor driven by an exonuclease (21) indicate that other enzymes can be used to construct artificial BBR motors. These studies suggest that BBR motors powered by enzymes with high turnover rates, not only natural ones but also engineered ones, could achieve fast chemical reaction cycles and short pause lengths, leading to the high speed.

Importantly, as shown in Fig. 8, such fast chemical reaction cycle leads to rotational diffusion-limited motion for micron-sized DNA-particle motors, where the stepping duration becomes comparable to or even longer than the pause length and limits the achievable speed. Fast motor proteins such as kinesin-1 and myosin V possess motor domains of ∼10 nm in size, which enable fast diffusion. Therefore, nano-sized DNA-particle motors will be essential to achieve the speed of several hundred nm s^−1^ to 1 µm s^−1^, which is comparable to those of the fast motor proteins.

### Improvement of upper limit of processivity and run-length associated with entrap detachment

Our results also clarify how processivity and run-length depend on the particle size and the balance of the kinetic parameters. The strong diameter dependence of the processivity and run-length (Fig. 4, Table 2) is explained by a change in the detachment mode, the stochastic detachment for small particles and low DNA/RNA hybridization rate, and the entrap detachment for large particles and/or high DNA/RNA hybridization rate (Fig. 4e, f). When the entrap detachment is major, the processivity reaches a maximum value around 1000 (Table 2), and the run-length gradually increases with the particle size due to the gradual increase in the step size (Fig. 3, Fig. 4g, h).

Although our simulation reproduced overall trend of particle diameter dependence of the processivity and run-length, it did not reach the very long run-length of ∼200 µm reported experimentally for 5000-nm DNA-silica microparticle motors (Table 1 and 2) (16). Interestingly, in the previous study, the 5000-nm particle motor resumed the motion even after apparent entrapment, without clear detachment from the surface. This behavior can be explained by slow Brownian motion and large mass of the micron-sized particle. Even if all DNA/RNA duplexes are hydrolyzed, the micron-sized particle will not diffuse away from the surface immediately, because of its slow Brownian motion and gravitational settling toward the surface. Instead, micron-sized particle will show quasi two-dimensional Brownian motion on the surface and easily rebind to the intact RNA sites. This will result in very long run-length. Note that in our study, each simulation is terminated when the number of DNA/RNA duplexes reaches zero and subsequent rebinding events are not considered.

In the two-dimensional BBR motors, only the backward step is prohibited but forward and side steps are allowed, and their motions cannot be perfectly unidirectional in principle (Fig. 5). Therefore, the upper limit of the processivity and run-length caused by the entrap detachment is determined by the degree of unidirectionality or unidirectionality index (Fig. 5b, Table 2). If the unidirectionality is further improved, the processivity and run-length will be further improved. For example, in the case of perfectly unidirectional ballistic motion (or in the case that only the forward steps are allowed), the entrap detachment will not occur, and very high processivity and very long run-length are possible (about the improvement of the unidirectionality, see next section).

Another possible strategy to suppress the entrap detachment is rail restoration, in which hydrolyzed (burnt) rails are replaced with intact ones by the strand displacement or repaired by enzymatic synthesis, so that the motor never becomes entrapped inside a fully burnt region. Alternative approach may be introducing additional, non-degrading binding sites, in which a subset of motor–surface interactions are just reversible and do not accompany enzymatic cleavages. It has been reported that influenza A viruses operate as a BBR motor on the receptor-modified surface, and employ neuraminidase-mediated receptor cleavage and hemagglutinin-mediated reversible receptor binding to achieve persistent motions (30–33). Analogously, the DNA-nano/microparticle motors with mixed degradative and non-degradative interactions could remain associated with the surface while slowing the buildup of the fully burnt regions, thereby extending the processivity and run-length.

### Improvement of unidirectionality: effect of particle size, multivalency, and anisotropic structures

Our analysis of step angles shows how unidirectionality is affected by the perfectness of the BBR mechanism and the multivalency of the interaction at the motor–surface interface (Fig. 5). Large particles exhibited step-angle distributions that are more tightly focused in the forward direction (Fig. 5a), and the unidirectionality index increased with the particle diameter (Fig. 5b, Table 2). RNA hydrolysis efficiency under the motor trajectory also increased with the particle diameter and positively correlates with the unidirectionality index (Fig. 5b, d), consistent with the BBR mechanism.

Our detailed analysis also revealed how high multivalency further amplifies the directional bias (Fig. 6). When a large number of DNA/RNA duplexes are formed, the overlap of the mobile regions of all bound RNA sites is statistically biased and sharpened toward the forward direction relative to the previous step (Fig. 6b). As a result, large particle exhibits a strong forward bias in the next step direction, leading to high unidirectionality. Therefore, micron-sized particle is better than nano-sized particle to achieve high unidirectionality. However, as discussed above, the micron-sized particle has a significant drawback in achieving high speed (Fig. 8).

As an alternative strategy, unidirectionality can be improved by introducing anisotropic structures in the motor bodies or rails as demonstrated in the previous studies (16,34). Anisotropic rod-shape DNA-motors or one-dimensional RNA surface rails will restrict the directions of rotational diffusion, bias the centroid of all bound RNA sites and the overlap of mobile regions of all bound RNA sites to the forward direction. To achieve very fast and very long unidirectional motions comparable to motor proteins, nano-sized anisotropic motor bodies can be prepared with gold nanorods (35) or DNA-origami rods (34), and one-dimensional long RNA surface rails can be prepared by micro/nanofabrication (16) or DNA nanotechnology (36–38). Our geometry-based kinetic simulations can be extended to the artificial BBR motors which possess the anisotropic structures, and our next goal will be the quantitative predictions and the experimental verifications of their motility and performance metrics.

## Data availability

The data are available from the corresponding author upon reasonable request. Modeling code and analysis scripts are available from the corresponding author upon request.

## Author contributions

T.H. and R.I. designed research; T.H. developed geometric-based kinetic simulation and analyzed data. T.H. and R.I. wrote the manuscript.

## Acknowledgments

This work was supported by JSPS KAKENHI, Grants-in-Aid for Transformative Research Areas, “Materials Science of Meso-Hierarchy” Grant Number JP24H01732 to T.H. and “Molecular Cybernetics” Grant Number JP23H04434 to R.I., Grant-in-Aid for Early-Career Scientists Grant Number JP23K13645 to T.H., JST, ACT-X Grant Number JPMJAX24LE, to T.H., the grant of OML Project by the National Institutes of Natural Sciences (NINS program No, OML022501) to T.H., and Tatematsu Foundation. to T.H.

## References

1. Finer, J. T., R. M. Simmons, and J. A. Spudich, 1994. Single myosin molecule mechanics: piconewton forces and nanometre steps. Nature 368:113–119, doi: 10.1038/368113a0.

2. Huxley, H. E., 1969. The mechanism of muscular contraction. Science 164:1356–1365, doi: 10.1126/science.164.3886.1356.

3. Svoboda, K., C. F. Schmidt, B. J. Schnapp, and S. M. Block, 1993. Direct observation of kinesin stepping by optical trapping interferometry. Nature 365:721–727, doi: 10.1038/365721a0.

4. Vale, R. D., 2003. The molecular motor toolbox for intracellular transport. Cell 112:467–480, doi: 10.1016/s0092-8674(03)00111-9.

5. Walker, J. E., 2013. The ATP synthase: the understood, the uncertain and the unknown. Biochem. Soc. Trans. 41:1–16, doi: 10.1042/BST20110773.

6. Iino, R., K. Kinbara, and Z. Bryant, 2020. Introduction: Molecular Motors. Chem. Rev. 120:1–4, doi: 10.1021/acs.chemrev.9b00819.

7. Lund, K., A. J. Manzo, N. Dabby, N. Michelotti, A. Johnson-Buck, J. Nangreave, S. Taylor, R. Pei, M. N. Stojanovic, N. G. Walter, E. Winfree, and H. Yan, 2010. Molecular robots guided by prescriptive landscapes. Nature 465:206–210, doi: 10.1038/nature09012.

8. Yin, P., H. Yan, X. G. Daniell, A. J. Turberfield, and J. H. Reif, 2004. A Unidirectional DNA Walker That Moves Autonomously along a Track. Angew. Chem. Int. Ed. 43:4906–4911, doi: 10.1002/anie.200460522.

9. Cha, T.-G., J. Pan, H. Chen, J. Salgado, X. Li, C. Mao, and J. H. Choi, 2013. A synthetic DNA motor that transports nanoparticles along carbon nanotubes. Nat. Nanotechnol. 9:39–43, doi: 10.1038/nnano.2013.257.

10. Pan, J., T.-G. Cha, F. Li, H. Chen, N. A. Bragg, and J. H. Choi, 2017. Visible/near-infrared subdiffraction imaging reveals the stochastic nature of DNA walkers. Sci. Adv. 3:e1601600, doi: 10.1126/sciadv.1601600.

11. Imai, H., T. Shima, K. Sutoh, M. L. Walker, P. J. Knight, T. Kon, and S. A. Burgess, 2015. Direct observation shows superposition and large scale flexibility within cytoplasmic dynein motors moving along microtubules. Nat. Commun. 6:8179, doi: 10.1038/ncomms9179.

12. Schnitzer, M. J., and S. M. Block, 1997. Kinesin hydrolyses one ATP per 8-nm step. Nature 388:386–390, doi: 10.1038/41111.

13. Sakamoto, T., I. Amitani, E. Yokota, and T. Ando, 2000. Direct Observation of Processive Movement by Individual Myosin V Molecules. Biochem. Biophys. Res. Commun. 272:586–590, doi: 10.1006/bbrc.2000.2819.

14. Nakamura, A., K. I. Okazaki, T. Furuta, M. Sakurai, and R. Iino, 2018. Processive chitinase is Brownian monorail operated by fast catalysis after peeling rail from crystalline chitin. Nat. Commun. 9:3814, doi: 10.1038/s41467-018-06362-3.

15. Piranej, S., L. Zhang, A. Bazrafshan, W. Deng, and K. Salaita, 2026. Programming DNA machines to move. Nat. Rev. Chem., doi: 10.1038/s41570-025-00791-7.

16. Yehl, K., A. Mugler, S. Vivek, Y. Liu, Y. Zhang, M. Fan, E. R. Weeks, and K. Salaita, 2016. High-speed DNA-based rolling motors powered by RNase H. Nat. Nanotechnol. 11:184–190, doi: 10.1038/nnano.2015.259.

17. Bazrafshan, A., M. E. Kyriazi, B. A. Holt, W. Deng, S. Piranej, H. Su, Y. Hu, A. H. El-Sagheer, T. Brown, G. A. Kwong, A. G. Kanaras, and K. Salaita, 2021. DNA Gold Nanoparticle Motors Demonstrate Processive Motion with Bursts of Speed Up to 50 nm Per Second. ACS Nano 15:8427–8438, doi: 10.1021/acsnano.0c10658.

18. Harashima, T., A. Otomo, and R. Iino, 2025. Rational engineering of DNA-nanoparticle motor with high speed and processivity comparable to motor proteins. Nat. Commun. 16:729, doi: 10.1038/s41467-025-56036-0.

19. Korosec, C. S., I. N. Unksov, P. Surendiran, R. Lyttleton, P. M. G. Curmi, C. N. Angstmann, R. Eichhorn, H. Linke, and N. R. Forde, 2024. Motility of an autonomous protein-based artificial motor that operates via a burnt-bridge principle. Nat. Commun. 15:1511, doi: 10.1038/s41467-024-45570-y.

20. Zhang, L., S. Piranej, A. Namazi, S. Narum, and K. Salaita, 2024. “Turbo-Charged” DNA Motors with Optimized Sequence Enable Single-Molecule Nucleic Acid Sensing. Angew. Chem. Int. Ed. 63:e202316851, doi: 10.1002/anie.202316851.

21. Imtiaz, Y., J. Hardin, B. Sheyitov, L. Zhang, A. Foote, M. H. B. A. Rahman, K. Jackson, S. Piranej, A. Bazrafshan, and K. Salaita, 2025. DNA Motors Powered by Exonuclease III for Autonomous Rolling Motion and Biosensing Applications. bioRxiv, doi: 10.1101/2025.11.26.690764 (preprint posted November 26, 2025).

22. Korosec, C. S., L. Jindal, M. Schneider, I. Calderon de la Barca, M. J. Zuckermann, N. R. Forde, and E. Emberly, 2021. Substrate stiffness tunes the dynamics of polyvalent rolling motors. Soft Matter 17:1468–1479, doi: 10.1039/d0sm01811b.

23. Blanchard, A. T., S. Piranej, V. Pan, and K. Salaita, 2022. Adhesive Dynamics Simulations of Highly Polyvalent DNA Motors. J. Phys. Chem. B 126:7495–7509, doi: 10.1021/acs.jpcb.2c01897.

24. Kerssemakers, J. W., E. L. Munteanu, L. Laan, T. L. Noetzel, M. E. Janson, and M. Dogterom, 2006. Assembly dynamics of microtubules at molecular resolution. Nature 442:709–712, doi: 10.1038/nature04928.

25. Isojima, H., R. Iino, Y. Niitani, H. Noji, and M. Tomishige, 2016. Direct observation of intermediate states during the stepping motion of kinesin-1. Nat. Chem. Biol. 12:290–297, doi: 10.1038/nchembio.2028.

26. Block, S. M., L. S. B. Goldstein, and B. J. Schnapp, 1990. Bead movement by single kinesin molecules studied with optical tweezers. Nature 348:348–352, doi: 10.1038/348348a0.

27. Thorn, K. S., J. A. Ubersax, and R. D. Vale, 2000. Engineering the Processive Run Length of the Kinesin Motor. J. Cell Biol. 151:1093–1100, doi: 10.1083/jcb.151.5.1093.

28. Nakamura, A., T. Tasaki, Y. Okuni, C. Song, K. Murata, T. Kozai, M. Hara, H. Sugimoto, K. Suzuki, T. Watanabe, T. Uchihashi, H. Noji, and R. Iino, 2018. Rate constants, processivity, and productive binding ratio of chitinase A revealed by single-molecule analysis. Phys. Chem. Chem. Phys. 20:3010–3018, doi: 10.1039/c7cp04606e.

29. Pierobon, P., S. Achouri, S. Courty, A. R. Dunn, J. A. Spudich, M. Dahan, and G. Cappello, 2009. Velocity, processivity, and individual steps of single myosin V molecules in live cells. Biophys. J. 96:4268–4275, doi: 10.1016/j.bpj.2009.02.045.

30. Hamming, P. H., N. J. Overeem, and J. Huskens, 2020. Influenza as a molecular walker. Chem. Sci. 11:27–36, doi: 10.1039/c9sc05149j.

31. Liu, M., E. de Vries, and C. A. M. de Haan, 2025. Virion motility of sialoglycan-cleaving respiratory viruses. npj Viruses 3:59, doi: 10.1038/s44298-025-00140-x.

32. Sakai, T., S. I. Nishimura, T. Naito, and M. Saito, 2017. Influenza A virus hemagglutinin and neuraminidase act as novel motile machinery. Sci. Rep. 7:45043, doi: 10.1038/srep45043.

33. Ziebert, F., and I. M. Kulić, 2021. How Influenza’s Spike Motor Works. Phys. Rev. Lett. 126:218101, doi: 10.1103/PhysRevLett.126.218101.

34. Bazrafshan, A., T. A. Meyer, H. Su, J. M. Brockman, A. T. Blanchard, S. Piranej, Y. Duan, Y. Ke, and K. Salaita, 2020. Tunable DNA Origami Motors Translocate Ballistically Over μm Distances at nm/s Speeds. Angew. Chem. Int. Ed. 59:9514–9521, doi: 10.1002/anie.201916281.

35. Zhou, C., X. Duan, and N. Liu, 2015. A plasmonic nanorod that walks on DNA origami. Nat. Commun. 6:8102, doi: 10.1038/ncomms9102.

36. Green, L. N., H. K. K. Subramanian, V. Mardanlou, J. Kim, R. F. Hariadi, and E. Franco, 2019. Autonomous dynamic control of DNA nanostructure self-assembly. Nat. Chem. 11:510–520, doi: 10.1038/s41557-019-0251-8.

37. Rothemund, P. W. K., A. Ekani-Nkodo, N. Papadakis, A. Kumar, D. K. Fygenson, and E. Winfree, 2004. Design and Characterization of Programmable DNA Nanotubes. J. Am. Chem. Soc. 126:16344–16352, doi: 10.1021/ja044319l.

38. Zhan, P., K. Jahnke, N. Liu, and K. Göpfrich, 2022. Functional DNA-based cytoskeletons for synthetic cells. Nat. Chem. 14:958–963, doi: 10.1038/s41557-022-00945-w.

