## Supporting Table S1, Fig. S1, Fig. S2 for "Artificial DNA-nano/microparticle motors: Factors governing speed, run-length, and unidirectionality revealed by geometry-based kinetic simulations"

Takanori Harashima[1,2], Ryota Iino[1,2]
[1]Institute for Molecular Science, National Institutes of Natural Sciences, Okazaki, Aichi, Japan
[2]Graduate Institute for Advanced Studies, SOKENDAI, Hayama, Kanagawa, Japan

**Table S1. Geometric and kinetic parameters used in simulations.**

| | Particle diameter (nm) | DNA surface density (molecules nm$^{-2}$) | RNA surface density (molecules nm$^{-2}$) | RNA square lattice unit size (pixels) | RNA square lattice unit size (nm) | Accessible radius $R_{acc}$ (nm) | Mobile radius $R_{mobile}$ (nm) | $k_{on}^{DNA/RNA}$ (s$^{-1}$) | $k_{on}^{E}$[RNase H] (s$^{-1}$) | $k_{cat}^{E}$ (s$^{-1}$) |
|---|---|---|---|---|---|---|---|---|---|---|
| Fig. 2a, b | 100 | 0.1 | 0.022 | 600 | 1897 | 28 | 25.2 | 0.8 | 7.2 | 3 |
| | 500 | | | 600 | 1897 | 60.8 | 56.2 | | | |
| | 1000 | | | 700 | 2213 | 85.7 | 79.5 | | | |
| | 5000 | | | 1200 | 3794 | 191 | 178 | | | |
| Fig. 3a | 100 | 0.1 | 0.022 | 600 | 1897 | 28 | 25.2 | 0.2 | 7.2 | 3 |
| | 500 | | | 600 | 1897 | 60.8 | 56.2 | | | |
| | 1000 | | | 700 | 2213 | 85.7 | 79.5 | | | |
| | 5000 | | | 1200 | 3794 | 191 | 178 | | | |
| Fig. 3b | 100 | 0.1 | 0.022 | 600 | 1897 | 28 | 25.2 | 0.8 | 7.2 | 3 |
| | 500 | | | 600 | 1897 | 60.8 | 56.2 | | | |
| | 1000 | | | 700 | 2213 | 85.7 | 79.5 | | | |
| | 5000 | | | 1200 | 3794 | 191 | 178 | | | |
| Fig. 4c, d | 100 | 0.1 | 0.022 | 600 | 1897 | 28 | 25.2 | 0.8 | 7.2 | 3 |
| Fig. 4e, g | 100 | 0.1 | 0.022 | 600 | 1897 | 28 | 25.2 | 0.2 | 7.2 | 3 |
| | 500 | | | 600 | 1897 | 60.8 | 56.2 | | | |
| | 1000 | | | 700 | 2213 | 85.7 | 79.5 | | | |
| | 5000 | | | 1200 | 3794 | 191 | 178 | | | |
| Fig. 4f, h | 100 | 0.1 | 0.022 | 600 | 1897 | 28 | 25.2 | 0.8 | 7.2 | 3 |
| | 500 | | | 600 | 1897 | 60.8 | 56.2 | | | |
| | 1000 | | | 700 | 2213 | 85.7 | 79.5 | | | |
| | 5000 | | | 1200 | 3794 | 191 | 178 | | | |
| Figs. 5, 6 | 100 | 0.1 | 0.022 | 600 | 1897 | 28 | 25.2 | 0.8 | 7.2 | 3 |
| | 500 | | | 600 | 1897 | 60.8 | 56.2 | | | |
| | 1000 | | | 700 | 2213 | 85.7 | 79.5 | | | |
| | 5000 | | | 1200 | 3794 | 191 | 178 | | | |

| | | | | | | | | | | |
|---|---|---|---|---|---|---|---|---|---|---|
| Fig. 7 Figs. S1, S2 | 100 | 0.1 | 0.022 | 600 | 1897 | 28 | 25.2 | 0.2 | 0.036, 0.144, 0.36, 0.72, 1.44, 3.6, 7.2, 14.4, 36, 72 | 3 |
| | | | | | | | | 0.8 | | 3 |
| | | | | | | | | 0.8 | | 12 |
| | | | | | | | | 0.8 | | 30 |
| | | | | | | | | 2 | | 3 |
| | | | | | | | | 2 | | 12 |
| | | | | | | | | 2 | | 30 |
| | | | | | | | | 8 | | 3 |
| | | | | | | | | 8 | | 12 |
| | | | | | | | | 8 | | 30 |
| Fig. 8b, d, f | 100 | 0.1 | 0.022 | 600 | 1897 | 28 | 25.2 | 0.8 | 7.2 | 3 |
| | 500 | | | 600 | 1897 | 60.8 | 56.2 | | | |
| | 1000 | | | 700 | 2213 | 85.7 | 79.5 | | | |
| | 5000 | | | 1200 | 3794 | 191 | 178 | | | |
| Fig. 8c, e, g | 100 | 0.1 | 0.022 | 600 | 1897 | 28 | 25.2 | 8 | 72 | 30 |
| | 500 | | | 600 | 1897 | 60.8 | 56.2 | | | |
| | 1000 | | | 700 | 2213 | 85.7 | 79.5 | | | |
| | 5000 | | | 1200 | 3794 | 191 | 178 | | | |

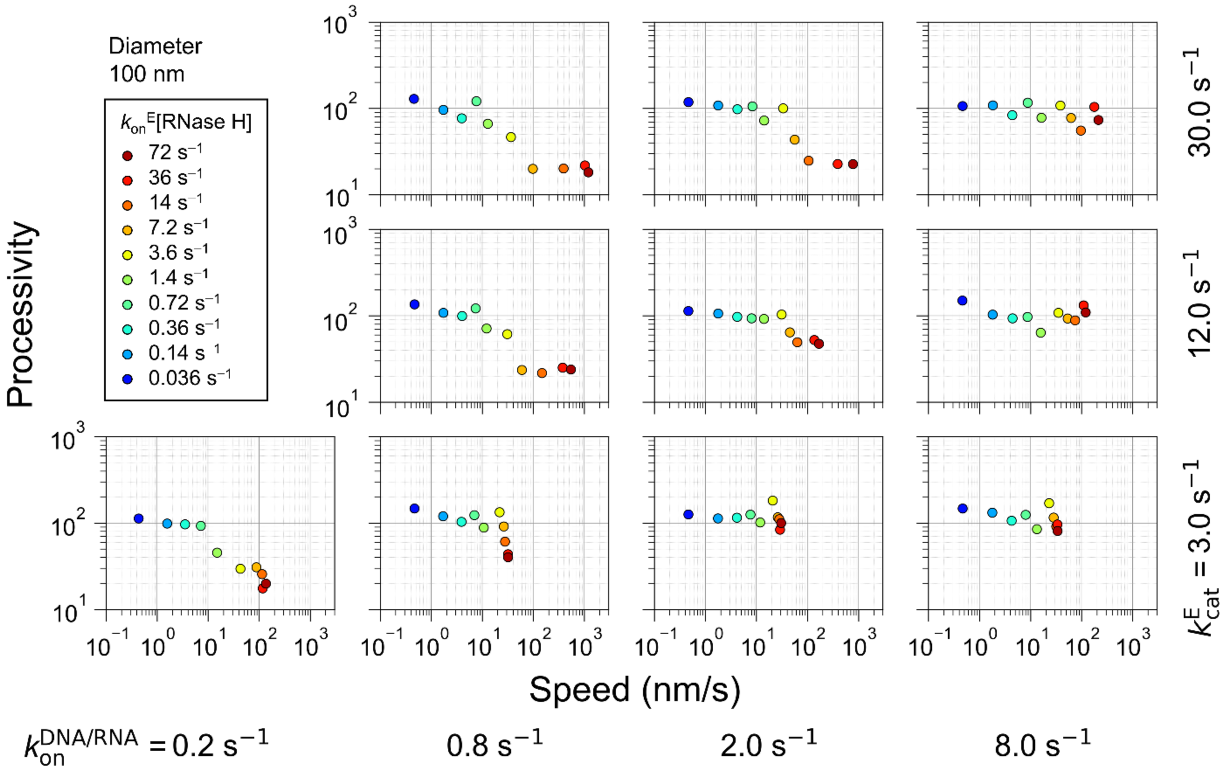


**Figure S1. Relationship between speed and processivity under various kinetic parameters for the motor with 100 nm particle.**

Scatter plots of median speed versus mean processivity obtained from 50 simulations of the motor with 100 nm particle under various kinetic conditions. Individual data points correspond to different combinations of DNA/RNA hybridization rates ($k_{on}^{DNA/RNA}$, from left to right panels), RNase H binding rates ($k_{on}^{E}$[RNase H], symbols with different colors in each panel), and RNA hydrolysis rates ($k_{cat}^{E}$, from bottom to top panels).

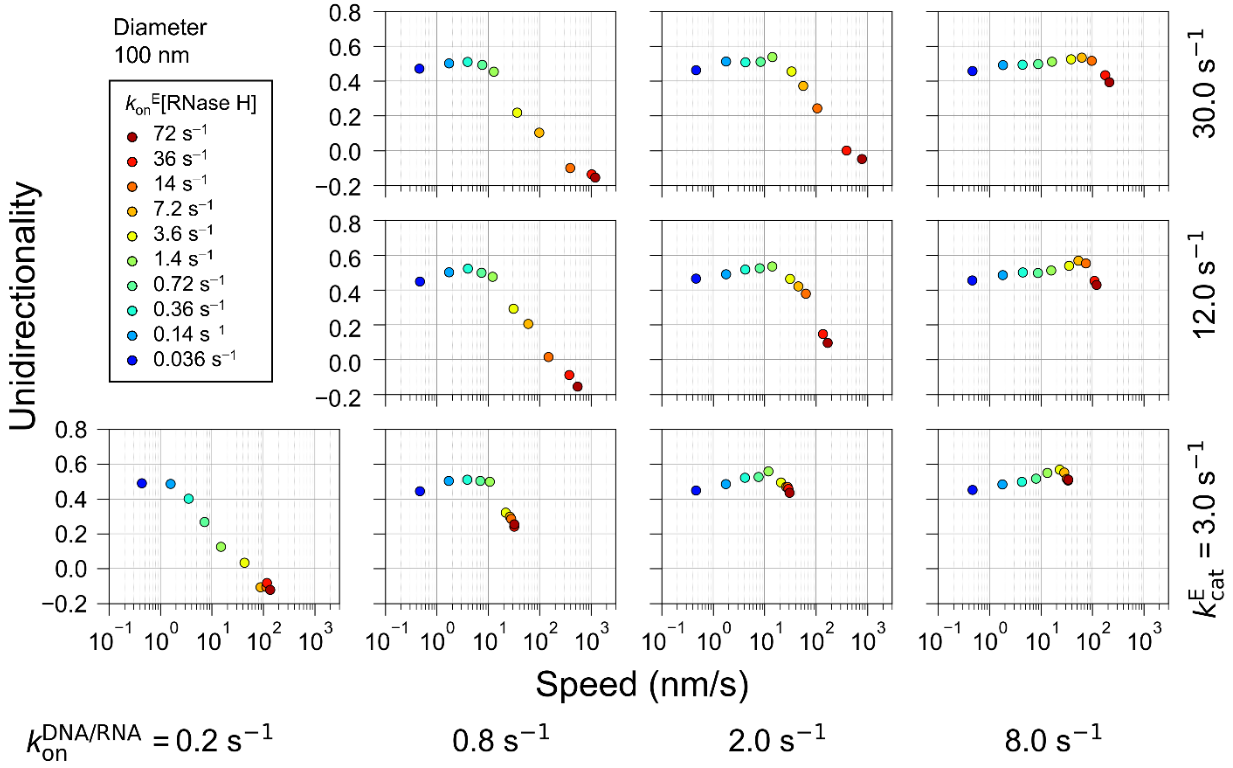


**Figure S2. Relationship between speed and unidirectionality under various kinetic parameters for the motor with 100 nm particle.**

Scatter plots of median speed versus unidirectionality index obtained from 50 simulations of the motor with 100 nm particle under various kinetic conditions. Individual data points correspond to different combinations of DNA/RNA hybridization rates ($k_{on}^{DNA/RNA}$, from left to right panels), RNase H binding rates ($k_{on}^{E}$[RNase H], symbols with different colors in each panel), and RNA hydrolysis rates ($k_{cat}^{E}$, from bottom to top panels).
